# Nirmatrelvir Resistant SARS-CoV-2 Variants with High Fitness in Vitro

**DOI:** 10.1101/2022.06.06.494921

**Authors:** Yuyong Zhou, Karen Anbro Gammeltoft, Line Abildgaard Ryberg, Long V. Pham, Ulrik Fahnøe, Alekxander Binderup, Carlos Rene Duarte Hernandez, Anna Offersgaard, Carlota Fernandez-Antunez, Günther Herbert Johannes Peters, Santseharay Ramirez, Jens Bukh, Judith Margarete Gottwein

**Affiliations:** Copenhagen Hepatitis C Program (CO-HEP), Department of Infectious Diseases, Copenhagen University Hospital–Hvidovre, 2650 Hvidovre, Denmark and Department of Immunology and Microbiology, Faculty of Health and Medical Sciences, University of Copenhagen, 2200 Copenhagen, Denmark; Department of Chemistry, Technical University of Denmark, 2800 Kongens Lyngby, Denmark

## Abstract

The oral protease inhibitor nirmatrelvir is expected to play a pivotal role for prevention of severe cases of coronavirus disease 2019 (COVID-19). To facilitate monitoring of potentially emerging resistance, we studied severe acute respiratory syndrome coronavirus 2 (SARS-CoV-2) escape from nirmatrelvir. Resistant variants selected in cell culture harbored different combinations of substitutions in the SARS-CoV-2 main protease (Mpro). Reverse genetic studies in a homologous infectious cell culture system revealed up to 80-fold resistance conferred by the combination of substitutions L50F and E166V. Resistant variants had high fitness increasing the likelihood of occurrence and spread of resistance. Molecular dynamics simulations revealed that E166V and L50F+E166V weakened nirmatrelvir-Mpro binding. The SARS-CoV-2 polymerase inhibitor remdesivir retained activity against nirmatrelvir resistant variants and combination of remdesivir and nirmatrelvir enhanced treatment efficacy compared to individual compounds. These findings have implications for monitoring and ensuring treatment programs with high efficacy against SARS-CoV-2 and potentially emerging coronaviruses.

---

The severe acute respiratory syndrome coronavirus 2 (SARS-CoV-2) protease inhibitor nirmatrelvir (PF-07321332) has recently been authorized as a first-line treatment for patients with a high risk for progression from mild-to-moderate to severe coronavirus disease 2019 (COVID-19)^1,2^. Moreover, nirmatrelvir is being considered for treatment of long term negative health effects of SARS-CoV-2 infection, termed long-COVID^3,4^. However, its high clinical efficacy^5^ might be threatened by development of antiviral resistance, as previously observed for virtually all small-molecule inhibitors targeting viruses such as influenza, hepatitis B and C, and human immune deficiency virus^6^.

We aimed to identify nirmatrelvir resistance associated substitutions and to investigate fitness of nirmatrelvir resistant variants. Moreover, we investigated sensitivity of these variants to the parenteral SARS-CoV-2 polymerase inhibitor remdesivir, approved for patients with severe COVID-19 or an increased risk for progression to severe COVID-19^7–9^, and evaluated prospects for combination of nirmatrelvir and remdesivir.

We previously characterized efficacy of the protease inhibitor boceprevir, structurally related to nirmatrelvir (Fig.S1), against SARS-CoV-2 in VeroE6 monkey kidney and A549-hACE2 human lung cells^10^. Here, we initially identified boceprevir resistance associated substitutions by induction of SARS-CoV-2 escape in VeroE6 cells (TableS1). While the original SARS-CoV-2 was suppressed by 3-fold 50% effective concentration (EC50) boceprevir, escape viruses could overcome up to 7-fold EC50 boceprevir; higher boceprevir concentrations could not be applied due to its relatively high cytotoxicity^10^. Polyclonal escape viruses from 4 independent escape experiments acquired substitutions L50F and A173V in the main viral protease Mpro (also termed nsp5 or 3CLpro) (TableS1). Compared to the original virus, selected polyclonal escape viruses, BOC-EV1 and BOC-EV2, showed up to 4.7-fold increased EC50 in short-term concentration-response treatments (Fig.1a), and resistance was confirmed in longer-term treatments (Fig.1b). When nirmatrelvir became available, we first confirmed its high efficacy against SARS-CoV-2, including similar efficacy against epidemiologically relevant variants, and its low cytotoxicity (Fig.1a; Fig.S2-4)^11–15^. The observed cell-line specific differences in EC50 are explained by differences in the expression level of efflux transporter P-glycoprotein^11,16^. Next, we demonstrated that polyclonal boceprevir escape viruses conferred up to 6.8-fold cross-resistance to nirmatrelvir in short-term concentration-response treatments (Fig.1a) and confirmed nirmatrelvir resistance in longer-term treatments (Fig.1b). We then induced SARS-CoV-2 escape from nirmatrelvir. Original SARS-CoV-2 was suppressed by 7-fold EC50 nirmatrelvir; however, nirmatrelvir escape viruses could overcome up to 40- and 120-fold EC50 nirmatrelvir in two different escape experiments, respectively (Fig.1c); higher nirmatrelvir concentrations inhibited spread of escape viruses. Polyclonal nirmatrelvir escape viruses harboring Mpro substitutions T21I+T304I (NIR-EV1) or L50F+E166V (NIR-EV2) conferred up to 5.9- or 175-fold nirmatrelvir resistance, respectively (Fig.1d, TableS2).

**Fig.1.**
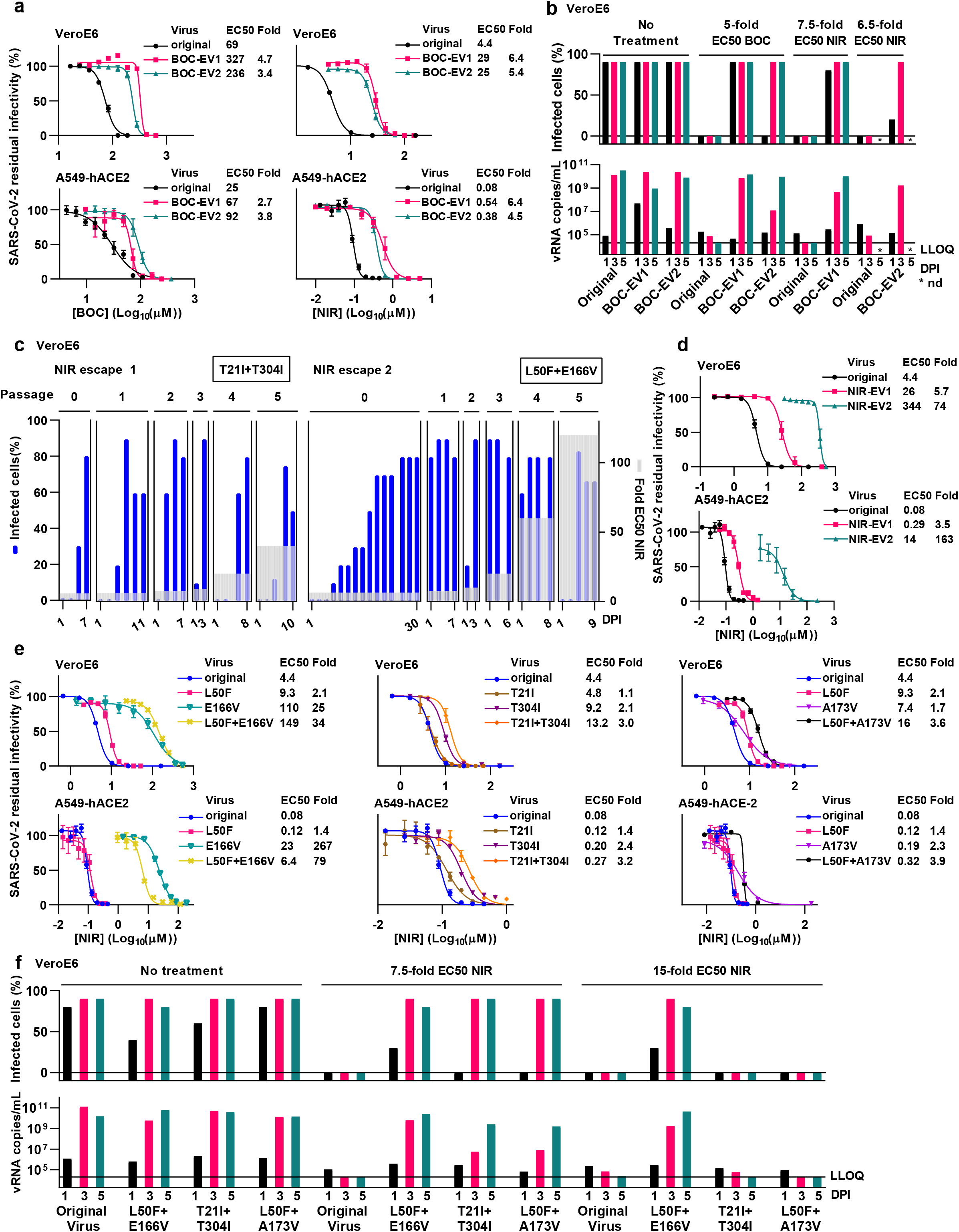
Identification of SARS-CoV-2 nirmatrelvir resistance associated substitutions in an infectious cell culture system. **(a)** Short-term concentration-response treatments of SARS-CoV-2 original virus and specified boceprevir escape viruses (BOC-EV1 and BOC-EV2) in VeroE6 or A549-hACE2 cells with boceprevir (BOC) or nirmatrelvir (NIR). Infected cells were visualized by spike protein immunostaining. Datapoints represent % residual infectivity, calculated by relating the number of infected cells in treated cultures to the mean number of infected cells in infected nontreated cultures, and are means of at least 4 replicates ± standard errors of the means (SEM). Curves and 50% effective concentrations (EC50) were determined in GraphPad Prism. Fold resistance (Fold) was determined as EC50_variant_/EC50_original virus_ using rounded values. **(b)** Longer-term treatments of SARS-CoV-2 original virus and specified boceprevir escape viruses (BOC-EV1 and BOC-EV2) in VeroE6 cells with the specified fold-EC50 of boceprevir (BOC) or nirmatrelvir (NIR). On the specified days post infection (DPI), the % SARS-CoV-2 infected culture cells were determined by spike protein immunostaining, and viral RNA (vRNA) titers in culture supernatants were determined by RT-qPCR. LLOQ, lower limit of quantification of the RT-qPCR assay. nd, not determined. **(c)** Induction of SARS-CoV-2 escape from nirmatrelvir (NIR) by serial viral passage under increasing drug pressure. Two escape experiments, each comprising a primary escape culture (passage 0) and 5 passage cultures (passage 1-5) were done; acquired Mpro substitutions are specified in boxes (compare TableS2). Left y-axes: % SARS-CoV-2 infected culture cells determined by spike protein immunostaining; right y-axes: fold-EC50 nirmatrelvir applied. X-axes: day post infection (DPI) for each culture. **(d)** Short-term concentration-response treatments of specified nirmatrelvir escape viruses (NIR-EV1 and NIR-EV2) in VeroE6 or A549-hACE2 cells with nirmatrelvir (NIR). For further details, see (a). **(e)** Short-term concentration-response treatments of variants with specified engineered resistance associated substitutions in VeroE6 or A549-hACE2 cells with nirmatrelvir (NIR). For further details, see (a). **(f)** Longer-term treatments of variants with specified engineered resistance associated substitutions in VeroE6 cells with the specified fold-EC50 nirmatrelvir (NIR). For further details, see (b).

We confirmed the significance of the identified putative resistance substitutions by reverse genetics using a bacterial artificial chromosome (BAC) clone corresponding to the SARS-CoV-2 variant used for escape experiments^17,18^. In short-term treatments, the L50F+E166V variant showed up to 80-fold nirmatrelvir resistance, with resistance being conferred by E166V; the T21I+T304I and L50F+A173V variants showed no-to-little resistance (Fig.1e). In longer-term treatments, all three double mutants spread under 7.5-fold EC50 nirmatrelvir, while only the L50F+E166V variant could spread under 15-fold EC50 nirmatrelvir (Fig.1f).

Engineered double mutants showed high fitness in transfection and passage cultures with infectivity titers comparable to those of the original virus (Fig.2a) and did not acquire additional substitutions in Mpro following 4 viral passages (TableS3). The fitness cost of single substitutions E166V and A173V was compensated by L50F (Fig.2a; Table.S3).

**Fig.2.**
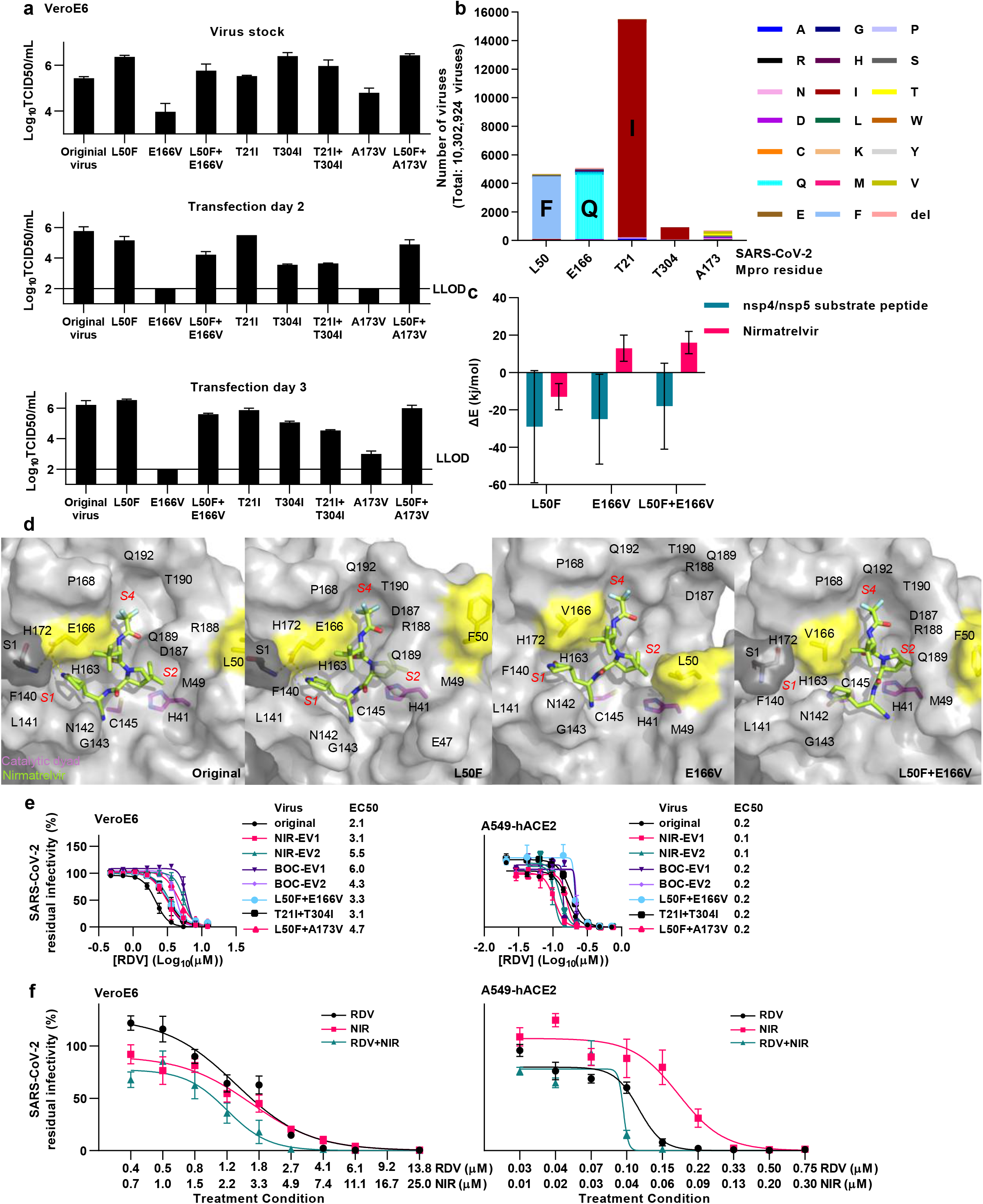
Identified SARS-CoV-2 resistant variants had high fitness, showed impaired molecular Mpro-nirmatrelvir interactions and retained sensitivity to remdesivir. **(a)** Viral infectivity titers of virus stocks generated from supernatants of viral passage cultures or of supernatants derived from transfected cultures for SARS-CoV-2 variants with specified engineered resistance associated substitutions in VeroE6 cells. Datapoints represent 50% tissue culture infectious doses (TCID50)/ml and are means of 4 replicates with SEM. LLOD, lower limit of detection. **(b)** Number of viruses identified in a GISAID-database search with specified amino acid substitutions at specified Mpro positions associated with nirmatrelvir resistance. del, deletion. **(c)** Interaction energy differences (ΔE_variant-original_) for binding of nirmatrelvir or of the nsp4/nsp5 substrate peptide to Mpro with L50F, E166V or E166V+L50F compared to original Mpro, as determined from MD simulations. **(d)** Final Mpro-nirmatrelvir conformations extracted from MD simulations with labelling of subsites *S1, S2* and *S4* (red), and subsite residues (black). The important interactions of E166 in monomer A (light grey) with S1 in monomer B (dark grey) and with nirmatrelvir (green) are indicated by yellow dashes. The catalytic dyad (H41 and C145) is shown in magenta and L50F and E166V in yellow. Non-carbon atoms are colored as follows, N: blue, O: red, F: light blue. **(e)** Short-term concentration-response treatments of specified boceprevir escape viruses (BOC-EV1 and BOC-EV2), nirmatrelvir escape viruses (NIR-EV1 and NIR-EV2) or variants with specified engineered resistance associated substitutions in VeroE6 or A549-hACE2 cells with remdesivir (RDV). For further details see Fig.1a. **(f)** Short-term concentration-response treatments of original SARS-CoV-2 virus in VeroE6 or A549-hACE2 cells with remdesivir (RDV), nirmatrelvir (NIR) or a combination of both (RDV+NIR). Each of the treatment condition (indicated by ticks on x-axis) was defined by the specified concentrations of RDV and NIR, which were applied singly and in combination, resulting in three datapoints per treatment condition. For further details see Fig.1a.

Analysis of sequences from the GISAID database (April 18th, 2022) revealed that resistance associated Mpro positions were overall conserved but permitted a certain degree of variation (Fig.2b, TableS4).

Mpro molecular dynamics simulations (Supporting Information Note 1 and Fig.S5) showed that E166V and L50F+E166V weakened and that L50F improved nirmatrelvir binding (Fig.2c, TableS5). In contrast, substrate binding was improved or unchanged for these substitutions (Fig.2c, TableS5), in line with previous observations for the single substitutions^19^. E166 has important roles in the coronavirus Mpro structure contributing to Mpro dimer stability through interactions with S1 in the other monomer^20^ and stabilizing substrate binding in a catalytically competent conformation through interactions in subsites *S1* and *S4^21^* (Fig.2d and Fig.S6). The fact that E166V weakened binding of Mpro to nirmatrelvir could be explained by loss of the enzymatically important^21^ nirmatrelvir-E166-S1 interactions and loss of nirmatrelvir interactions with Mpro residues 187-192 leading to an opening of subsites *S2* and *S4* (Fig.2d and Fig.S9-S10). L50F likely improved Mpro-nirmatrelvir binding by a gain of nirmatrelvir interactions with R188 and T190 at subsite *S4* (Fig.2d and Fig.S9-S10). In the double mutant, in addition to this gain of interactions, a reduction of nirmatrelvir interactions with S1, F140, L141, H163 and H172 in subsite *S1* was observed (Fig.2d and Fig.S9-S10). Consequently, E166V and L50F+E166V might shift the nirmatrelvir-Mpro conformational equilibrium towards non-catalytically competent states, reducing the probability of nirmatrelvir inhibition by 50% and 63%, respectively (Fig.S11). The weakened nirmatrelvir binding and reduced inhibition probability likely explain the experimentally observed resistance for E166V and L50F+E166V.

Finally, we demonstrated that remdesivir showed similar efficacy against nirmatrelvir resistant variants and the original SARS-CoV-2 (Fig.2e). Combination treatment with nirmatrelvir and remdesivir showed enhanced efficacy compared to treatment with the individual compounds, with residual infectivity in combination treatments being up to 5.7- and 3.9-fold decreased compared to treatments with nirmatrelvir and remdesivir, respectively (Fig.2f).

In conclusion, we report the identification of substitutions in the SARS-CoV-2 main protease conferring significant resistance to the first-in-class oral protease inhibitor nirmatrelvir, which is currently rolled-out globally. In clinical trials, nirmatrelvir maximum serum concentrations (Cmax) of 4.4 μM were reported^2^. Thus, Cmax/EC50 ratios of 1 and 55 for the original SARS-CoV-2 were reduced to 0.03 and 0.7 for the most resistant L50F+E166V variant and to 0.01 and 0.3 for the most resistant polyclonal escape virus in VeroE6 and A549-hACE2 cells, respectively. Resistance selection and spread in populations might be supported by the high fitness of resistant variants, a certain flexibility of resistance associated Mpro residues and suboptimal treatment adherence during home medication. Sensitivity of nirmatrelvir resistant variants to remdesivir and enhanced efficacy of combination treatments might provide an incentive for evaluation of nirmatrelvir-remdesivir combination therapy in clinical trials. Overall, this study provides a basis for monitoring of nirmatrelvir resistance substitutions in populations. As nirmatrelvir showed broad activity against different coronaviruses^16^, our findings do not only have implications for the current but also for future coronavirus outbreaks.

## Acknowledgements

We thank Dr. Bjarne Ørskov Lindhardt (Copenhagen University Hospital-Hvidovre) and Prof. Charlotte Menne Bonefeld (University of Copenhagen) for support, as well as Lotte Mikkelsen and Anna-Louise Sørensen (Copenhagen University Hospital-Hvidovre) for laboratory assistance. We thank Prof. Jean Dubuisson and Dr. Sandrine Belouzard for providing VeroE6 cells.

## Author Contributions

Study design: Y.Z., K.A.G., L.A.R, G.H.J.P., J.B., J.M.G. Acquisition of data: Y.Z., K.A.G., L.A.R., L.P., U.F., A.M.B., C.R.D.H., A.O., C.F.-A. Analysis and interpretation of data: Y.Z., K.A.G., L.A.R., U.F., A.M.B., C.R.D.H., G.H.J.P., S.R., J.B., J.M.G. Drafting of the manuscript: Y.Z., K.A.G., L.A.R., G.H.J.P., J.M.G. All authors reviewed the manuscript. Study supervision: J.M.G.

## Online Methods

### Cell culture

African green monkey kidney VeroE6 cells (gift from J. Dubuisson) were cultured in Dulbecco’s Modified Eagle Medium (Invitrogen, Paisley, UK) as described^10,22,23^. Human lung epithelial A549 cells expressing human angiotensin converting enzyme 2 receptor (hACE2), referred to as A549-hACE2 cells, (Invivogen, Toulouse, France) were cultured in Dulbecco’s Modified Eagle Medium: Nutrient Mixture F-12 (Gibco, Paisley, UK) as described^10,22,23^. Both culture media were supplemented with 10% heat inactivated fetal bovine serum (Sigma, Saint Louis Missouri, USA), 100 U/mL penicillin and 100μg/mL streptomycin (Gibco/Invitrogen corporation, Carlsbad, California, USA). A549-hACE2 culture medium was in addition supplemented with 0.5 μg/mL puromycin (Invivogen, Toulouse, France). Cells were maintained at 37°C and 5% CO_2_ and subcultured every 2-3 days.

### SARS-CoV-2 virus stocks

All virus stocks were generated in VeroE6 cells and sequence confirmed by next generation sequencing (NGS) in relation to the patient isolate or BAC clone they originated from as described in the section *Next generation sequencing of SARS-CoV-2 genomes*. SARS-CoV-2 original virus: SARS-CoV-2 D614G. The 2^nd^ viral passage stock with an infectivity titer of 5.5 log_10_ 50% tissue culture infectious doses (TCID50)/mL was based on the isolate SARS-CoV-2/human/Denmark/DK-AHH1/2020^17^ (GenBank: MZ049597), with a spike protein sequence as reported in early outbreak isolates such as the Wuhan-Hu-1 isolate with the D614G mutation. This stock was used for induction of viral escape and as a reference in short-term concentration-response treatments and longer-term treatments.

SARS-CoV-2 alpha variant. The 2^nd^ viral passage stock with an infectivity titer of 6.7 log_10_ TCID50/mL was based on the isolate SARS-CoV-2/human/DNK/DK-AHH1/2021 (Genbank: OK041529)^24^. This stock was used for short-term concentration-response treatments. SARS-CoV-2 delta variant. The 2^nd^ viral passage stock with an infectivity titer of 6.0 log_10_ TCID50/mL was based on the isolate SARS-CoV-2/human/DNK/DK-AHH3/2021 (Genbank: in submission). This stock was used for short-term concentration-response treatments.

SARS-CoV-2 omicron A1 variant. The 2^nd^ viral passage stock with an infectivity titer of 5.5 log_10_ TCID50/mL was based on the isolate SARS-CoV-2/human/DNK/DK-AHH4/2021 (Genbank in submission). This stock was used for short-term concentration-response treatments.

Polyclonal escape virus stocks were generated and used as described in the section *Induction of SARS-CoV-2 escape*.

Stocks of SARS-CoV-2 recombinants were generated and used as described in the section *SARS-CoV-2 recombinants and transfections*.

### SARS-CoV-2 recombinants and transfections

Recombinants were based on the BAC clone encoding the sequence of the SARS-CoV-2 D614G isolate SARS-CoV-2/human/Denmark/DK-AHH1/2020^18^. For generation of variants, point mutations were engineered by infusion and mega-primer based cloning using sequence specific primers.

In vitro RNA transcripts reflecting the SARS-CoV-2 genome were generated as described^18^. In brief, NotI linearized plasmids were purified using the Zymo DNA clean&concentrator-25 kit with Zymo-Spin™ IC-XL columns (ZR BAC DNA Miniprep Kit, ZymoResearch, Irvine, CA, USA) and subjected to in vitro transcription with the mMESSAGE mMACHINE T7 Transcription Kit (ThermoFisher, Waltham, MA,USA). Transfection of 200,000 VeroE6 cells per well plated the previous day in 12-well plates (Thermo Fisher Scientific, Roskilde, Denmark) was carried out using 2 μg of RNA transcripts, quantified with the Qubit RNA BR Assay Kit (ThermoFisher, Waltham, MA, USA) and lipofectamine2000 (ThermoFisher) in Opti-MEM (Invitrogen, Waltham, MA, USA); as an exception, for the L50F, A173V and L50F+A173V variants, 1 μg of RNA transcripts were used. Cell culture supernatants were collected on selected days post transfection and stored at −80°C. Transfection efficacy was evaluated by inspection of transfection cultures for cytopathogenic effects (CPE) in an inverted light microscope and determination of supernatant SARS-CoV-2 infectivity titers as described in the section *Determination of SARS-CoV-2 infectivity titers*.

1^st^ viral passage virus stocks were generated by inoculation of VeroE6 cells, plated the previous day at 3 million cells in T80 flasks (Thermo Fisher Scientific, Roskilde, Denmark), with 250 μl supernatant derived from transfection cultures at the expected peak of infection at day 2 or 3 post transfection as confirmed by observation of CPE in an inverted light microscope. As an exception for the L50F and L50F+A173V variants a 2^nd^ passage was done for virus stock generation. From infected 1^st^ or 2^nd^ passage cultures cell culture supernatants were collected at the peak of infection, subjected to NGS for sequence confirmation and stored at −80°C. Virus stocks were used for short-term concentration-response treatments and longer-term treatments.

### Serial passage of SARS-CoV-2 variants

To investigate their genetic stability SARS-CoV-2 variants were subjected to four viral passages. VeroE6 cells, plated the previous day at 1 million cells in T25 flasks (Thermo Fisher Scientific, Roskilde, Denmark), were inoculated with 250 μl supernatant derived at the peak of infection from the preceding culture. Supernatant derived at the peak of infection from the 4^th^ passage culture was subjected to NGS.

### Inhibitors

All inhibitors were purchased from Acme Bioscience (Palo Alto, California, USA) and dissolved in DMSO (Sigma, Saint Louis, Missouri, USA).

### Cell viability assays

To evaluate cytotoxic effects of the studied inhibitors, cell viability assays were done as described^10,22,23^ using the CellTiter 96 AQueous One Solution Cell Proliferation Assay (Promega, Madison, WI, USA). VeroE6 or A549-hACE2 cells, plated the previous day at 10,000 cells per well in 96-well flat-bottom plates (Thermo Fischer Scientific, Roskilde, Denmark), were treated with the specified concentrations of inhibitors, including 3 replicate wells per concentration and 10 nontreated control wells. Following incubation for 46-50 hours at 37°C and 5% CO_2_, CellTiter 96 AQueous One Solution Reagent was added at 20 μL per well followed by incubation for 1.5 to 2 h at 37°C and 5% CO_2_ and determination of absorbance at 492 nm using a FLUOstar OPTIMA 96-well plate reader (BMG LABTECH, Offenburg, Germany). For calculation of cell viability in %, absorbance values from individual treated wells were related to the mean absorbance of nontreated control wells. Datapoints are means of triplicates with SEM. Sigmoidal concentration-response curves were fitted and 50% cytotoxic concentration (CC50) values calculated using GraphPad Prism 8.0.0 with a bottom constraint of 0 applying the equation Y= Top/(1+10^(Log10EC50-X)*HillSlope^).

### Induction of SARS-CoV-2 escape

VeroE6 cells, plated the previous day at 1 million cells in T25 flasks, were inoculated at 0.00002 multiplicity of infection (MOI) with the SARS-CoV-2 D614G stock and treated with boceprevir or nirmatrelvir at concentrations selected to suppress but not eradicate viral infection. Every 2-3 days, cell culture supernatants were collected and stored at - 80°C for NGS, cells were subcultured as required aiming to maintain semi-confluent cell layers and fresh supernatant containing inhibitors at specified concentrations was added. Upon subculturing replicate cultures were plated on chamber slides for immunostainings to determine the percentage of SARS-CoV-2 infected culture cells as described in the section *SARS-CoV-2 spike protein immunostaining*. Results from immunostainings were used to adapt inhibitor concentrations. For boceprevir, 5 escape experiments were carried out, further detailed in TableS1. For nirmatrelvir, 2 escape experiments were carried out, each comprising a primary escape culture followed by 5 passages under increasing inhibitor concentrations, further detailed in TableS2.

Boceprevir polyclonal escape virus 1 and 2 (BOC-EV1 and BOC-EV2) stocks were generated by inoculation of VeroE6 cells, plated the previous day at 3 million cells in T80 flasks, with 15 μl supernatant derived on day 74 and 57 from escape 1 and 2 (TableS1), respectively. Nirmatrelvir polyclonal escape virus 1 and 2 (NIR-EV1 and NIR-EV2) stocks were generated by inoculation of VeroE6 cells, plated at 3 million cells the previous day in T80 flasks, with 15 μl supernatant derived from passage 5 day 10 from escape 1 or passage 5 day 9 from escape 2. From these inoculated cultures, supernatants were collected at the peak of infection, stored at −80°C and subjected to NGS. Virus stocks were used for short-term concentration-response treatments and longer-term treatments.

### SARS-CoV-2 spike protein immunostaining

Immunostaining was used for monitoring of SARS-CoV-2 infected VeroE6 cell cultures for induction of viral escape and longer-term treatments as described^10,23^. In these cultures, following cell splitting, replicate cultures were plated in 8-well chamber slides (Thermo Fisher Scientific, Rochester, NY, USA). The next day, slides were submerged in methanol (J.T.Baker, Gliwice, Poland) for 20 minutes to fix cells and inactivate SARS-CoV-2. Slides were washed 3 times with PBS (Sigma, Gillingham, UK) with 0.1% Tween-20 (Sigma, Saint Louis, Missouri) (PBS-tween) and incubated with 1^st^ antibody SARS-CoV-2 spike chimeric monoclonal antibody (Sino Biological #40150-D004, Beijing, China) diluted 1:1,000 in PBS with 1% bovine serum albumin (Roche, Mannheim, Germany) and 0.2% skim milk (Easis, Aarhus, Denmark) (PBSK) for 2 hours. Slides were washed 2 times with PBS-tween and incubated with 2^nd^ antibody Alexa-Fluor 488 goat anti-human IgG (H+L) (Invitrogen #A-11013, Paisley, UK) diluted 1:500 and Hoechst 33342 (Invitrogen, Paisley, UK) diluted 1:1,000 in PBS-tween for 20 minutes. Percentages of SARS-CoV-2 spike protein positive cells were evaluated by fluorescence microscopy (ZEISS Axio Vert.A1, Jena, Germany), using the following designations: 0% infected cells (no cells infected), single infected cells, and 10%–90% infected cells (in steps of 10%).

### Short-term concentration-response treatments

VeroE6 or A549-hACE2 cells, plated the previous day at 10,000 cells per well in 96-well flat-bottom plates (Thermo Fischer Scientific, Roskilde, Denmark), were inoculated with the specified SARS-CoV-2 virus stocks as described^10,22,23^; the size of the inoculum was determined in pilot assays aiming at 800-3000 infected cells in infected nontreated wells and absence of CPE at the end of the experiment. Cultures were incubated for 1 hour at 37°C and 5% CO_2_ and treated with specified concentrations of inhibitors as described^10^ with at least 4 replicates per concentration; at least 4 infected and nontreated and 8 noninfected and nontreated replicate cultures were included in each assay. Following incubation for 46-50 hours at 37°C and 5% CO_2_, plates were subjected to SARS-CoV-2 spike protein immunostaining. Plates were submerged in methanol for 20 minutes and washed 3 times with PBS-tween, followed by blocking of endogenous peroxidase activity with H_2_O_2_ for 10 minutes, 2 washes with PBS-tween and blocking by PBSK for 30 minutes. Plates were incubated with 1^st^ antibody SARS-CoV-2 spike chimeric monoclonal antibody (Sino Biological #40150-D004, Beijing, China) diluted 1:5,000 in PBSK for 2 hours, washed 2 times with PBS-tween and incubated for 1 hour with 2^nd^ antibody F(ab’)2-Goat anti-human IgG Fc Cross-Adsorbed Secondary Antibody, HRP (Invitrogen#A24476, Carlsbad, CA, USA) or Goat F(ab’)2 Anti-Human IgG – Fc (HRP), preadsorbed (Abcamab#98595, Cambridge, UK), diluted 1:2,000 in PBSK. Following 2 washes with PBS-tween, cells were stained with DAB substrate BrightDAB kit (Immunologic # BS04-110, Duiven, Netherlands).

The number of single SARS-CoV-2 spike protein positive cells per well was evaluated using an ImmunoSpot series 5 UV Analyzer (CTL Europe GmbH, Bonn, Germany). Representative images from concentration-response antiviral treatment assays are shown in Zhou and Gilmore et al^22^. Mean counts of noninfected nontreated wells, which were usually <100, were subtracted from counts of individual infected wells. For calculation of % residual infectivity counts of individual infected treated wells were related to mean counts of infected nontreated wells. Datapoints are means of at least 4 replicates with SEM. Sigmoidal concentration-response curves were fitted and EC50 values calculated as described previously using Graphpad Prism 8.0.0 with a bottom constraint of 0 applying the equation Y= Top/(1+10^(Log10EC50-X)*HillSlope^). Fold resistance (Fold) was determined as EC50_variant_/EC50_original virus_. EC50_original virus_ was the mean of several EC50 determinations shown in Fig.S2. In Fig.1a, d and e, a representative treatment curve is shown for the original SARS-Cov-2 virus and treatment curves included in the same graph were not in all instances carried out in the same experimental setup. However, in each treatment experiment with SARS-CoV-2 variants the original SARS-CoV-2 virus was included for comparison to ensure reproducibility.

### Longer-term treatments

VeroE6 cells, plated the previous day at 1 million cells in T25 flasks, were infected at 0.00002 MOI with the specified SARS-CoV-2 virus stocks as described^10^. Cells were treated with specified fold EC50 of inhibitors upon inoculation and then every 2 days upon subculturing. Upon subculturing, replicate cultures were plated on chamber slides for immunostainings to determine the percentage of SARS-CoV-2 infected cells as described in the section *SARS-CoV-2 spike protein immunostaining*. Further, cell culture supernatants were collected and stored at −80°C to determine viral RNA (vRNA) titers by RT-qPCR. An infected nontreated culture was included as a positive infection control.

### Combination treatments

Combination of nirmatrelvir with remdesivir for inhibition of SARS-CoV-2 was evaluated in short-term treatments in VeroE6 or A549-hACE2 cells as described^10^. VeroE6 or A549-hACE2 cells, plated the previous day at 10,000 cells per well in 96-well flat-bottom plates, were inoculated with the original SARS-CoV-2 virus stock; the size of the inoculum was determined in pilot assays aiming at 2000-4000 infected cells in infected nontreated wells and absence of CPE at the end of the experiment. Cultures were incubated for 1 hour at 37°C and 5% CO_2_ and treated with specified concentrations of inhibitors. For both inhibitors and the combination of dilution series were used spanning the inhibitor EC50 values and aiming at residual infectivity values between 0 and 100 %. In combination treatments the same concentrations as in single treatments were used with a fixed ratio. All treatment conditions were evaluated using at least 4 replicates including at least 8 infected nontreated replicates and 8 noninfected nontreated replicates. Following incubation for 46-50 hours at 37°C and 5% CO_2_, plates were subjected to SARS-CoV-2 spike protein immunostaining, automated counting of single SARS-CoV-2 spike protein positive cells, and further evaluation as described in the section *Short-term concentration-response treatments*. Fold-enhancement of combination treatments compared to treatments with individual compounds was calculated per treatment condition, which was defined by a specific concentration of nirmatrelvir and remdesivir, relating % residual infectivities to each other.

### Determination of SARS-CoV-2 RNA titers

The vRNA titers in cell culture supernatants were determined by qRT-PCR as described^10^. In brief, RNA was extracted using Trizol LS (Life Technologies) and chloroform (Sigma, Saint Louis, Missouri, USA) and purified using the Zymo RNA Clean and Concentrator-5 kit For RT-qPCR reactions the TaqMan Fast Virus 1-Step Master Mix (Thermo Fischer) was used with previously described primers and probes^25^: E_Sarbeco F (5’-ACAGGTACGTTAATAGTTAATAGCGT-3’), E_Sarbeco_R (5’-ATATTGCAGCAGTACGCACACA-3’) and E_Sarbeco_P (FAM-5’-ACACTAGCCATCCTTACTGCGCTTCG-3’-BHQ1) as well as the LifeCycler 96 System (Roche). The lower limit of quantification (LLOQ) of the assay was calculated as (mean of RNA titers in supernatants derived from noninfected control cultures) + (3 standard deviations). **Determination of SARS-CoV-2 infectivity titers**. SARS-CoV-2 infectivity titers in cell culture supernatants from transfection experiments were determined as described previously^26^. VeroE6 cells, plated the previous day at 10,000 cells per well in 96-well flat-bottom plates, were inoculated with culture supernatants in 10-fold dilution series. Following incubation for 70-74 hours at 37°C and 5% CO_2_, plates were subjected to SARS-CoV-2 spike protein immunostaining as described in the section *Short-term concentration-response treatments*. Plates were imaged using an ImmunoSpot series 5 UV Analyzer (CTL Europe GmbH) and scored infected or noninfected. Infectious titers were calculated as TCID50/ml using the Reed-Muench method. The LLOD was 2 Log_10_ TCID50/mL, defined by the used starting dilution.

### Next generation sequencing of SARS-CoV-2 genomes

SARS-CoV-2 RNA was extracted and purified as described in the section *Determination of SARS-CoV-2 RNA titers*. RT-PCR was used to generate five overlapping amplicons and NEBNext Ultra II FS DNA library Prep kit (New England BioLabs, Ipswich, Massachusetts, USA) was used for library preparations. NGS analysis was done as described^17,18,23,26^.

### Molecular dynamics simulations

Preparation of systems: Coordinates of SARS-CoV-2 Mpro with nirmatrelvir (Protein Data Bank (PDB) entry: 7vh8^27^) and nsp4/nsp5 substrate peptide (PDB entry: 7mgs^28^) bound, respectively, were obtained from the Protein Data Bank (PDB; Fig.S5). PyMOL^29^ version 2.5.0 was used for preparing the systems for simulation: building missing residues, deleting ions and ligands (other than nirmatrelvir), applying symmetry operations to form the Mpro dimer, and introducing substitutions (L50F, E166V or L50F+E166V). Furthermore, the covalent bond between C145 and nirmatrelvir was broken in the Mpro-nirmatrelvir structure and the intact nirmatrelvir nitrile warhead was built; further A145 in the Mpro-substrate peptide structure was mutated to C145 (Fig.S6). Protonation states for titratable residues at pH 7.4 were assigned based on *pK_a_* calculations using the H++ server^30^ and histidine protonation site was determined by analysis of the hydrogen bonding network around histidine. The N- and C-termini of the nsp4/nsp5 substrate peptide were kept neutral to avoid introduction of charges that are not present in the SARS-CoV-2 polyprotein nsp4-nsp5. The Mpro dimers were simulated in water boxes having a dodecahedron shape, and a minimum distance between solute and box edges of 12 Å was applied. Periodic boundary conditions were applied in all 3 Cartesian directions. Sodium and chloride ions were added to a concentration of 0.150 M NaCl corresponding to the cytosolic physiological ionic strength^31^. The CHARMM36m all-atom protein force field^32,33^ was used in the simulations, and a nirmatrelvir topology file was generated using the CHARMM General Force Field (CGenFF)^34,35^ using the CGenFF interface (https://cgenff.umaryland.edu).

#### Simulation settings

Simulations of Mpro dimers (original, L50F, E166V and L50F+E166V) with substrate peptide or nirmatrelvir molecules bound were run in triplicate using Gromacs version 2021.2^36,37^. The systems were minimized using a steepest descent algorithm until the maximum force was below 1000 kJ/mol/nm. The simulations were carried out using a leap-frog integrator and an integration time step of 1 fs. After minimization, an *NVT* equilibration step was carried out for 100 ps, followed by an *NPT* equilibration step for 100 ps. In both steps, position restraints were applied for Mpro-substrate peptide or Mpro-nirmatrelvir, and the temperature coupling was controlled out using a modified Berendsen thermostat^38^ with a time constant of 0.1 ps. For the *NPT* equilibration step, Berendsen pressure coupling^38^ was used with a time constant of 2.0 ps. The simulations were carried out in an *NPT* ensemble at 310 K and 1 bar for 100 ns, of which the first 50 ns were considered equilibration and the last 50 ns were the production run and used in the subsequent analysis. A modified Berendsen thermostat^38^ was used for temperature coupling with a time constant of 0.1 ps and the Parinello-Rahman approach^39,40^ was used for pressure coupling with a time constant of 2.0 ps. The Verlet cutoff scheme was used for calculating short-range van der Waals interactions with a cutoff at 12 Å in combination with force-switching starting at 10 Å. The Particle Mesh Ewald method^41,42^ was used for calculating long-range electrostatics with a 1.6 Å grid spacing. Hydrogen bonds were kept rigid using the LINCS algorithm^43^.

#### Analysis of simulations

Root-mean-square deviation (RMSD) of atom coordinates were calculated for all Mpro-nirmatrelvir (Fig.S7) or Mpro-substrate peptide (Fig.S8) atoms relative to the minimized structures using the gmx rms tool in Gromacs.

Interaction energies (E) between the Mpro dimer and substrate or nirmatrelvir, respectively, were extracted from the simulations using the gmx energy tool in Gromacs. The interaction energies calculated for each of the 2 substrate/nirmatrelvir molecules bound in the Mpro dimer were averaged, and the values in TableS5 and Fig.2c are averages of the dimer average interaction energies with SEM given as uncertainties. The differences in interaction energy between original and variant Mpro are calculated as: Δ*E* = *E_variant_* — *E_original_*. Thus, negative energy differences indicate improved nirmatrelvir/substrate binding to an Mpro variant, while positive differences indicate weakened binding.

Distance calculations were carried out using VMD version 1.9.4^44^. For the inhibition probability plots, distances were calculated between C145 sulfur and the nirmatrelvir cyano carbon, and between Gly143 amide nitrogen and the nirmatrelvir cyano nitrogen. For the interaction heat maps, the frequencies of nirmatrelvir-Mpro residues within 3 Å proximity were calculated and converted to percentage of the simulation frames the interactions occurred in. Differences in frequencies (Fig.S11) between original and variant Mpro were calculated as: Δfrequency(interactions) = *frequency*(*interactions*)*_variant_* – *frequency* (*interactions*) *_original_*. Only interactions with differences larger than 5% were included in Fig. S11.

## References

1. https://www.ema.europa.eu/en/news/covid-19-ema-recommends-conditional-marketing-authorisation-paxlovid.

2. https://www.fda.gov/media/155050/download.

3. Peluso, M. J. et al. (2022) doi:10.21203/RS.3.RS-1617822/V1.

4. Geng, L. N. et al. (2022) doi:10.21203/rs.3.rs-1443341/v1.

5. Hammond, J. et al. N. Engl. J. Med. 386, 1397–1408 (2022).

6. Mason, S. et al. Antiviral Res. 158, 103–112 (2018).

7. https://www.ema.europa.eu/en/medicines/human/EPAR/veklury.

8. https://www.samc.com/assets/documents/covid19/nursing/remdesivir_eua-hcp-fact-sheet-8-2020.pdf.

9. Beigel, J. H. et al. N. Engl. J. Med. 383, 1813–1826 (2020).

10. Gammeltoft, K. A. et al. Antimicrob. Agents Chemother. 65, (2021).

11. Rai, D. K. et al. (2021) doi:10.1101/2022.01.17.476644.

12. Vangeel, L. et al. Antiviral Res. 198, (2022).

13. Ullrich, S. et al. Bioorg. Med. Chem. Lett. 62, 128629 (2022).

14. Li, P. et al. Cell Res. 32, 322 (2022).

15. Abdelnabi, R. et al. Nat. Commun. 13, (2022).

16. Owen, D. R. et al. Science 374, 1586–1593 (2021).

17. Ramirez, S. et al. Antimicrob. Agents Chemother. 65, (2021).

18. Fahnøe, U. et al. Viruses 14, (2022).

19. Flynn, J. M. et al. bioRxiv 2022.01.26.477860 (2022).

20. Cheng, S. C. et al. Biophys. J. 98, 1327–1336 (2010).

21. Tan, J. et al. J. Mol. Biol. 354, 25–40 (2005).

## References

22. Zhou, Y. et al. Sci. Rep. 11, (2021).

23. Zhou, Y. et al. Viruses 13, (2021).

24. Sølund, C. et al. Vaccines 10, (2022).

25. Corman, V. M. et al. Eurosurveillance 25, 2000045 (2020).

26. Offersgaard, A. et al. Vaccines 9, (2021).

27. Zhao, Y. et al. Protein Cell (2021) doi:10.1007/S13238-021-00883-2.

28. MacDonald, E. A. et al. ACS Infect. Dis. 7, 2591–2595 (2021).

29. The PyMOL Molecular Graphics System, Version 2.5.0, Schrödinger, LLC.

30. Anandakrishnan, R. et al. Nucleic Acids Res. 40, (2012).

31. Cortese, J. D. et al. Biochim. Biophys. Acta 1100, 189–197 (1992).

32. Best, R. B. et al. J. Chem. Theory Comput. 8, 3257–3273 (2012).

33. Huang, J. et al. Nat. Methods 14, 71–73 (2017).

34. Vanommeslaeghe, K. et al. J. Comput. Chem. 31, 671–690 (2010).

35. Yu, W. et al. J. Comput. Chem. 33, 2451–2468 (2012).

36. Van Der Spoel, D. et al. J. Comput. Chem. 26, 1701–1718 (2005).

37. Abraham, M. J. et al. SoftwareX 1–2, 19–25 (2015).

38. Berendsen, H. J. C. et al. J. Chem. Phys. 81, 3684 (1998).

39. Parrinello, M. et al. J. Appl. Phys. 52, 7182 (1998).

40. Nosé, S. et al. http://dx.doi.org/10.1080/00268978300102851 50, 1055–1076 (2006).

41. Essmann, U. et al. J. Chem. Phys. 103, 8577 (1998).

42. Darden, T. et al. J. Chem. Phys. 98, 10089 (1998).

43. Hess, B. et al. J Comput Chem 18, 14631472 (1997).

44. Humphrey, W. et al. J. Mol. Graph. 14, 33–38 (1996).

