## Supporting Information for "Nirmatrelvir Resistant SARS-CoV-2 Variants with High Fitness in Vitro"

**Supplementary text**

**Note 1: Analysis of main protease (Mpro) molecular dynamics simulations**

***Mpro structure, function and inhibition***

Mpro (nsp5) is one of two cysteine proteases of severe acute respiratory syndrome coronavirus 2 (SARS-CoV-2). The function of Mpro is to cleave the SARS-CoV-2 polyproteins pp1a and pp1ab into mature nonstructural proteins at 11 cleavage sites[1]. The crystal structure of Mpro with a nsp4/nsp5 peptide bound[2] shown in Fig. a illustrates the nomenclature of the cleavage junction with P5-P1 corresponding to the five C-terminal residues of nsp4 and P1'-P2' corresponding to the two N-terminal residues of nsp5. The cleavage occurs between P1 and P1' (Fig. b). Nirmatrelvir is a covalent inhibitor of the Mpro, which has been designed to mimic substrate residues P4-P1 (Fig. c and d)[3].

Structures of Mpro with substrate peptides bound suggest that peptide bond cleavage is initiated by a nucleophilic attack on the carbonyl carbon (Fig. b) by the C145 thiolate, which is stabilized by the adjacent H41[4]. The attack leads to formation of a high-energy transition state with a negatively charged carbonyl oxygen[5], which is stabilized by hydrogen bonds to mainchain amide hydrogens of G143, S144 and C145 forming the oxyanion hole (Fig. a)[2].

For nirmatrelvir, a nucleophilic attack on the nitrile warhead by the C145 thiolate leads to a reversible covalent bond to C145 and thus Mpro inhibition[6]. Similarly to peptide bond breakage in natural substrates, the nucleophilic attack leads to high-energy transition states with a negatively charged nitrile nitrogen[7], which is stabilized in the structure by hydrogen bond interactions in the oxyanion hole with mainchain amide hydrogens of G143, S144 and C145 (Fig. b).

***Nirmatrelvir resistance associated substitutions***

In cell culture experiments, Mpro substitutions E166V and L50F+E166V are found to confer a high level of resistance to nirmatrelvir. E166 is a key residue in Mpro connecting the substrate binding site with the dimer interface[8]. In the substrate binding site, E166 stabilizes substrate binding by mainchain interactions with P3 and side chain interactions to the amide nitrogen in the P1 glutamine (Fig. a). These interactions are maintained when nirmatrelvir binds (Fig. b). E166 furthermore interacts with the N-terminal S1 in the other Mpro monomer, and is involved in substrate-induced Mpro dimerization[8]. As the Mpro monomers are not catalytically active, dimerization is essential for Mpro function[9].

Additionally, the S1-E166 interaction is essential for maintaining the correct shape of subsite *S1*[1]. This is supported by several studies of highly similar SARS-CoV Mpro variants. In one study, it was found that deletion of the N-terminal residues 1-3 in the SARS-CoV Mpro reduced the catalytic efficiency to 76% compared to the original Mpro and led to a slight reduction in dimerization[10]. In comparison, deletion of residues 1-4 led to a dramatic shift of the Mpro monomer-dimer equilibrium resulting in the monomer being the major species and led to a reduction of the catalytic efficiency to 1%[10]. Another study found that the mutation S1A did not affect the monomer-dimer equilibrium and resulted in 46% catalytic efficiency compared to original Mpro[11]. A third study found that E166A slightly reduced Mpro dimerization and reduced catalytic efficiency to 31% compared to the original Mpro. In combination with another mutation in the dimer interface, R298A, E166A shifted the Mpro monomer-dimer equilibrium towards monomer as the major species[8].

The results in these studies highlight the important role of the S1-E166-substrate interaction for maintaining Mpro enzymatic activity. While the S1-E166 intermonomer interaction contributes to the Mpro dimer stability, the interaction does not play a vital role for dimerization. The reduced enzymatic activity upon breakage of the S1-E166-substrate interaction is suggested to disturb the correct catalytically competent conformation of the substrate binding site including the oxyanion hole[12].

***Trajectory analysis for simulations***

The root-mean-square deviation (RMSD) evolutions of the nirmatrelvir and nsp4/nsp5 substrate peptide extracted from the simulations are shown in Fig. **S** and Fig, respectively. The RMSD values fluctuate around average values ranging from 1.8-2.4 Å indicating that the Mpro dimer is stable throughout the simulations and that the simulations have converged within 50 ns. We used the last 50 ns of the simulations for further analyses.

***Nirmatrelvir-Mpro interactions***

To evaluate how the interactions between nirmatrelvir and the SARS-CoV-2 Mpro changed for the Mpro variants, the changes in interaction patterns are illustrated in the heatmap in Fig. , and Mpro residues for which increased/decreased interactions are observed are shown in Fig and Fig.2d. In the following, the interaction patterns observed for L50F, E166V and L50F+E166V variants are discussed.

The L50F variant leads to an increase in interactions of R188 and T190 (Fig. ; 15% and 14%, respectively) with the P4 trifluoroacetyl group of nirmatrelvir (Figb and Fig.2d), and a reduction in interactions of M49 (Fig. ; -13%) with the P2 dimethylcyclopropylproline group of nirmatrelvir (Figb and Fig.2d).

The E166V substitution leads to a reduction of intermonomer interactions of S1 (Fig. ; -17%) with the P1 γ-lactam ring of nirmatrelvir (Figc and Fig.2d), and a reduction of interactions of D187 and Q192 (Fig. ; -13% and -15%, respectively) with the P2 dimethylcyclopropylproline group of nirmatrelvir leading to an opening of the binding pocket at subsite *S2* and *S4* as the loop formed by residues 187-192 moves away from nirmatrelvir (Figc and Fig.2d). We furthermore observe a gain of interactions with L50 (Fig. ; 11%) and the P2 dimethylcyclopropylproline group of nirmatrelvir (Figc and Fig.2d).

The double substitution L50F+E166V leads to a combination of the changes in interactions observed for the individual substitutions. In comparison with E166V, we observe a more dramatic reduction of intermonomer interactions of S1 (Fig. ; -32%) with the P1 γ-lactam ring of nirmatrelvir (Figd and Fig.2d). As for L50F, we observe an increase in interactions of R188 and T190 (Fig. ; 27% and 14%, respectively) with the P4 trifluoroacetyl group of nirmatrelvir (Figd and Fig.2d). These changes in interactions lead to a different binding pose of nirmatrelvir in the binding site, in which the P1 γ-lactam ring moves out of subsite *S1* (formed by F140, L141, H163, V166, H172 in one monomer (monomer A) and S1 in the other monomer (monomer B)), and the nitrile group moves towards T25, L27 and M49 (Figd and Fig.2d). The reductions of interactions of nirmatrelvir with F140, L141, H163, H172 are -17%, -15%, -14%, and -32%, respectively, and the gains of interactions of nirmatrelvir with T25, L27 and M49 are 11%, 10%, and 12%, respectively.

***Interaction energy analysis***

The interaction energies extracted from Mpro-nirmatrelvir and Mpro-substrate peptide MD simulations are presented in TableS5, and interaction energy differences are presented in Fig.2c. The relatively large errors for the substrate peptide simulations reflect the flexibility of the nsp4/nsp5 peptide.

For Mpro with E166V or L50F+E166V, we observe less favourable interaction energies for nirmatrelvir relative to original Mpro resulting mainly from reduction in the coulombic contribution to the interaction energies. This may be explained by the loss of S1-E166-nirmatrelvir interaction (Fig. ). For L50F and L50F+E166V variants, we observe slightly more favourable Lennard-Jones (LJ) energy contributions, which may be explained by a gain in interactions of nirmatrelvir with R188 and T190 (Fig).

***Inhibition probabilities***

For nirmatrelvir to successfully inhibit the SARS-CoV-2 Mpro, nirmatrelvir must bind to the Mpro in a conformation that is compatible with catalysis and thus leads to covalent attachment of nirmatrelvir to Mpro C145. We define a catalytically competent state by two criteria, 1) the sulfur atom in C145 and cyano carbon atom in nirmatrelvir should be in close enough proximity for the nucleophilic attack[7], and 2) the backbone amide of G143 in the oxyanion hole and cyano nitrogen atom in nirmatrelvir should be in close enough proximity for stabilizing the nirmatrelvir transition state[12]. The distances are shown in the Mpro structure in Fig. , and the numerical values extracted from the simulations are shown in Fig. . In agreement with other Mpro-substrate peptide/Mpro-nirmatrelvir simulation studies[7,13], we observe two populations of C145 sulfur and nirmatrelvir cyano carbon distances for the original Mpro. In 24% of the simulation frames, C145 sulfur and nirmatrelvir cyano carbon are in closer proximity than 4.3 Å, and in 76% of the simulation frames further apart than 4.3 Å. In all simulation frames, G143 amide nitrogen and nirmatrelvir cyano nitrogen are in closer proximity than 4.3 Å. Based on these findings, we consider conformations with C145 sulfur-nirmatrelvir cyano carbon distances below 4.3 Å and G143 amide nitrogen and nirmatrelvir cyano nitrogen distances below 4.3 Å as catalytically competent.

For the L50F variant, there is a slight decrease in the probability of nirmatrelvir-Mpro being in a catalytically competent conformation relative to original Mpro (24% to 20%). The probability for E166V to be in a catalytically competent state is reduced to 50% compared to the original Mpro (from 24% to 12%), while a third population with larger G143 amide nitrogen-nirmatrelvir cyano nitrogen distances is present for the L50F+E166V variant thus reducing the probability of a catalytically competent state to 63% compared to the original Mpro (from 24% to 15%).

***Conclusion***

In conclusion, we have found that the E166V and L50F+E166V variants gave rise to weakened Mpro-nirmatrelvir binding, whereas the L50F variant gave rise to improved Mpro-nirmatrelvir binding (TableS5 and Fig.2c). On the structural level, the weakened binding is explained by a loss of the enzymatically important S1-E166-nirmatrelvir interaction (Fig. , Fig and Fig.2d). For the E166V variant, this loss of interaction was accompanied by a loss of interaction in subsite *S2* (Fig. ) leading to an opening of the binding pocket (Figc and Fig.2d) and larger distances between the catalytic C145 and the nirmatrelvir cyano group reducing the probability of inhibition by 50 % compared to original Mpro (Fig. ).

For the L50F+E166V variant, the loss of S1-E166-nirmatrelvir interactions and gain in R188 and T190 interactions (Fig. ) led to a conformational change of the nirmatrelvir binding pose in which the P1 γ-lactam ring moved out of subsite *S1* (Figd and Fig.2d). Consequently, the distance between C143 in the oxyanion hole and the nirmatrelvir cyano group became too large for catalysis to take place reducing the probability of inhibition by 63% compared to original Mpro (Fig. ).

The above analyses of MD simulations provide an atomic-level explanation of the observed resistance against nirmatrelvir observed for the E166V and L50F+E166V variants in cell culture.

### References


1. Zhang, L. *et al. Science* **368**, 409–412 (2020).
2. MacDonald, E. A. *et al. ACS Infect. Dis.* **7**, 2591–2595 (2021).
3. Mótyán, J. A. *et al. Int. J. Mol. Sci.* **23**, (2022).
4. Lee, J. *et al. Nat. Commun. 2020 111* **11**, 1–9 (2020).
5. Świderek, K. *et al. Chem. Sci.* **11**, 10626–10630 (2020).
6. Zhao, Y. *et al. Protein Cell* (2021) doi:10.1007/S13238-021-00883-2.
7. Ngo, S. T. *et al. RSC Adv.* **12**, 3729–3737 (2022).
8. Cheng, S. C. *et al. Biophys. J.* **98**, 1327–1336 (2010).
9. Fan, K. *et al. J. Biol. Chem.* **279**, 1637–1642 (2004).
10. Hsu, W. C. *et al. J. Biol. Chem.* **280**, 22741–22748 (2005).
11. Chen, S. *et al. J. Biochem.* **143**, 525–536 (2008).
12. Tan, J. *et al. J. Mol. Biol.* **354**, 25–40 (2005).
13. Díaz, N. *et al. Chem. Commun. (Camb).* **57**, 5314–5317 (2021).

### Supplementary Figures

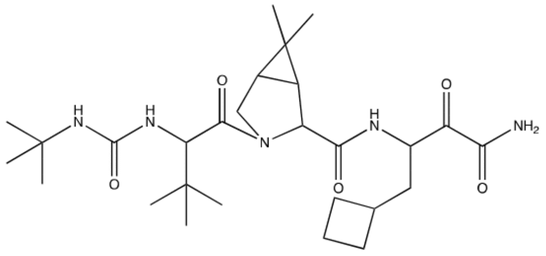

Boceprevir

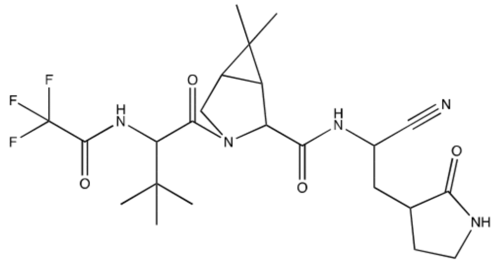

Nirmatrelvir

**Fig. S1**. **Structural formulas of boceprevir and nirmatrelvir.**

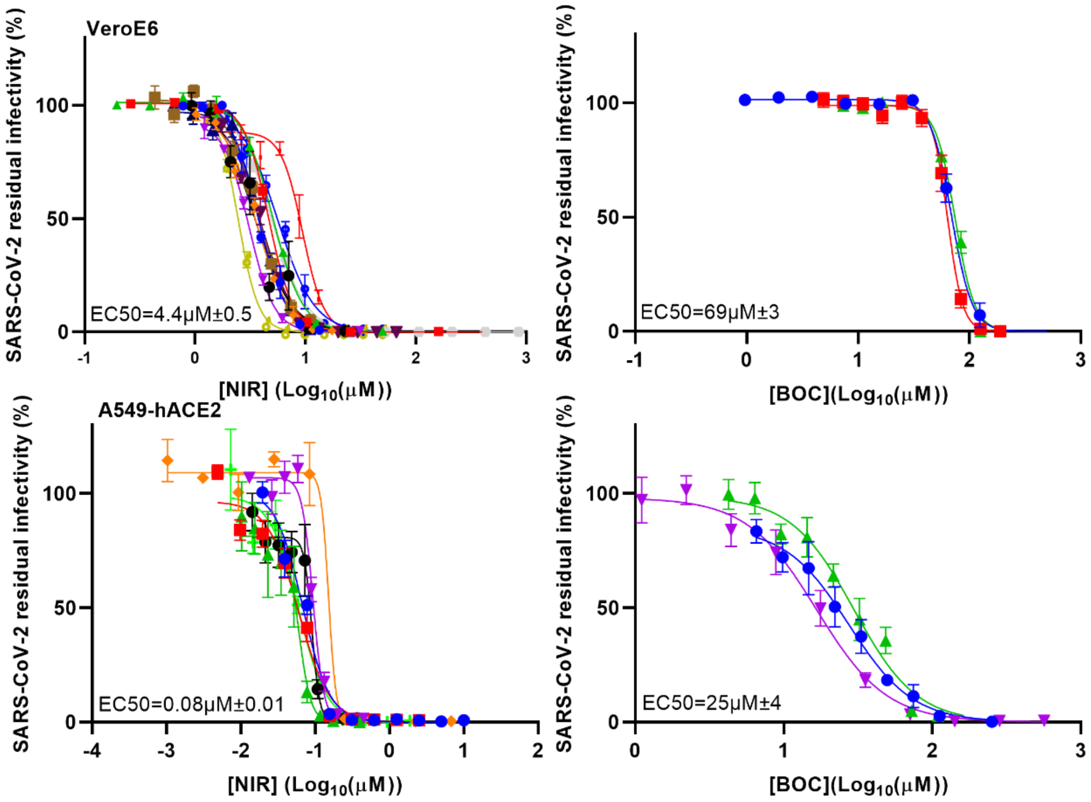


**Fig. S2. Determination of EC50 of boceprevir and nirmatrelvir against SARS-CoV-2 original virus in short-term concentration-response assays.** Short-term concentration-response treatments of SARS-CoV-2 original virus with boceprevir (BOC) or nirmatrelvir (NIR) in VeroE6 or A549-hACE2 cells in 96-well plates. Graphs show all treatments carried out during this study at different timepoints; treatments are color-coded; 13 and 7 nirmatrelvir treatments were done in VeroE6 and A549-hACE2 cells, respectively; 3 boceprevir treatments were done in VeroE6 and A549-hACE2 cells, respectively. Infected cells were visualized by spike protein immunostaining. Datapoints represent % residual infectivity, calculated by relating the number of infected cells in treated cultures to the mean number of infected cells in at least 8 infected nontreated cultures, and are means of at least 4 replicates ± standard errors of the means (SEM). Curves and 50% effective concentrations (EC50) were determined as described in Online Methods section *Short-term concentration-response treatments*. Given EC50 are the mean of the EC50 of individual curves with SEM and are used as reference EC50 for fold-resistance determinations.

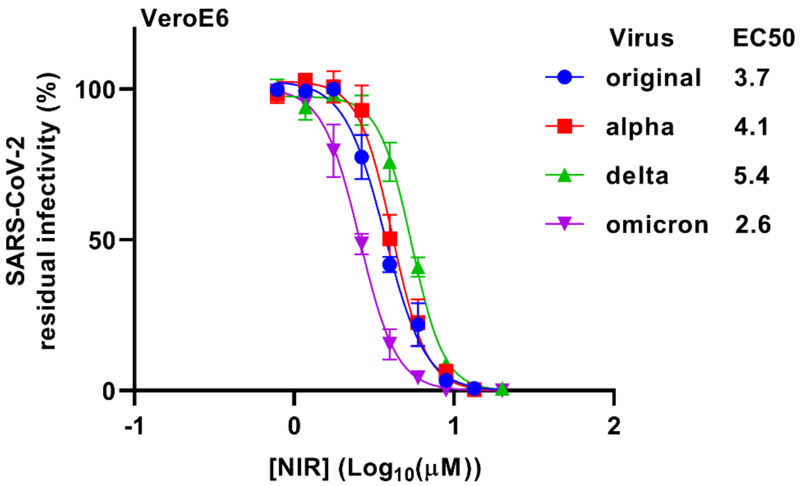


**Fig. S3. Nirmatrelvir had similar efficacy against the SARS-CoV-2 original virus as well as the alpha, delta and omicron variants.** Short-term concentration-response treatments of the SARS-CoV-2 original virus, as well as the alpha, delta and omicron variants with nirmatrelvir (NIR) were carried out in VeroE6 cells in 96-well plates. Infected cells were visualized by spike protein immunostaining. Datapoints represent % residual infectivity, calculated by relating the number of infected cells in treated cultures to the mean number of infected cells in at least 8 infected nontreated cultures, and are means of at least 4 replicates ± SEM. Curves and EC50 were determined as described in Online Methods *Short-term concentration-response treatments*.

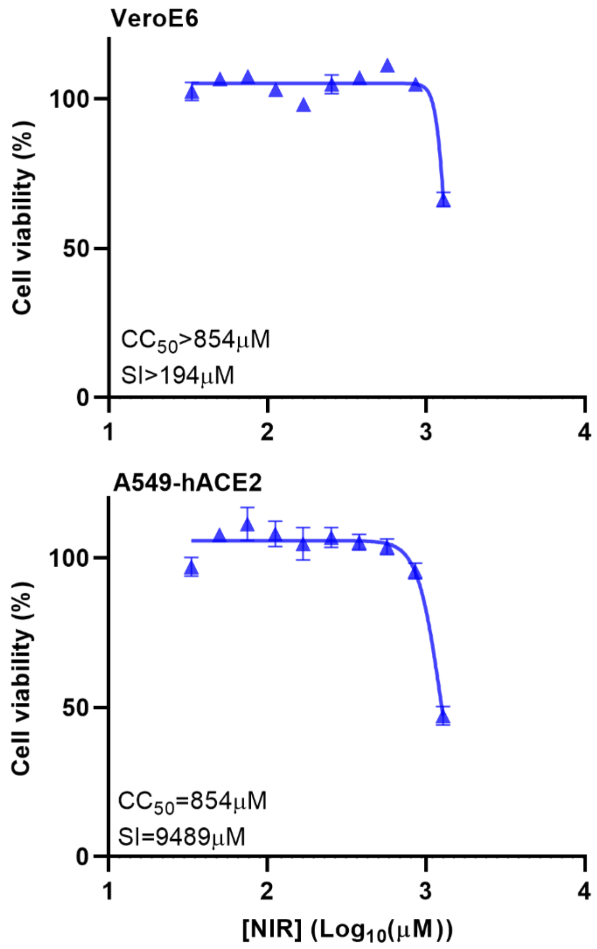


**Fig. S4. Nirmatrelvir had a low cytotoxicity and a high selectivity index.** Cell viability assays were carried out for nirmatrelvir (NIR) in VeroE6 and A549-hACE2 cells in 96-well plates. Datapoints represent % cell viability, calculated by relating the OD of treated cultures to the mean OD of 10 nontreated cultures, and are means of 3 replicates ± SEM. Curves and 50% cytotoxic concentrations (CC50) were determined using Graphpad Prism 8.0.0 applying the equation $Y= Top/(1+10^{(Log10EC50-X)*HillSlope})$ with a bottom constraint of 0. Selectivity indexes (SI) were determined as CC50/EC50.

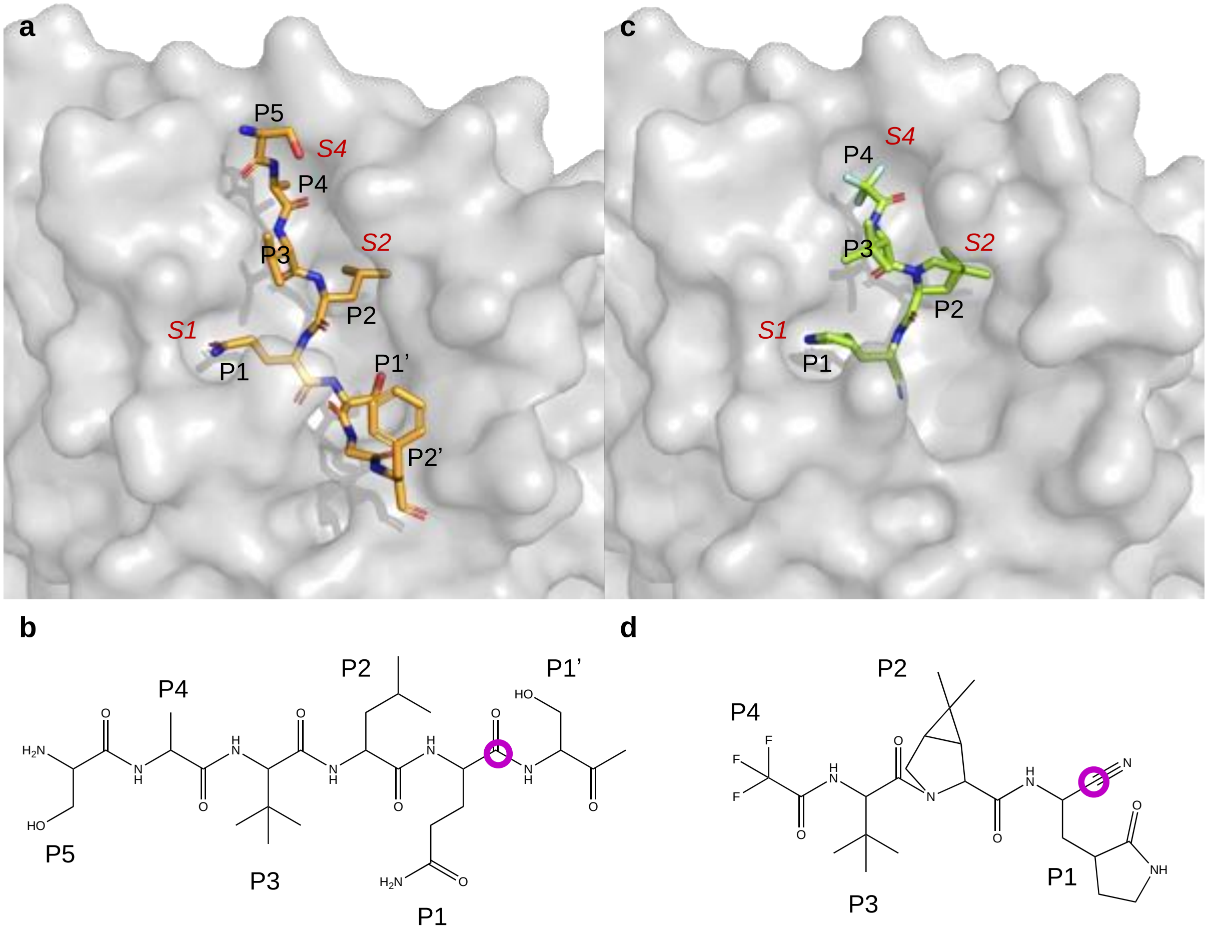


**Fig. S5. Structures of Mpro, nsp4-nsp5 substrate and nirmatrelvir. (a)** Crystal structure of Mpro (nsp5) with nsp4/nsp5 substrate peptide (orange sticks) bound with residues P5-P1 and P1'-P2' (PDB entry: 7mgs[2]). **(b)** Chemical structure of the nsp4/nsp5 junction with residues P5-P1 and P1'. The carbonyl carbon that is the target of the nucleophilic attack by C145 is highlighted with a purple circle. **(c)** Crystal structure of nsp5 with nirmatrelvir (green sticks) covalently bound with labelling of the structural elements mimicking substrate residues P4-P1 (PDB entry: 7vh8[6]). The Mpro is shown in a grey surface representation with subsites S1, S2, and S4 labelled in red and substrate / nirmatrelvir in stick modus with C: orange/green, N: blue, O:red, F:light-blue in (a) and (c). **(d)** Chemical structure of nirmatrelvir with labelling of the chemical groups mimicking substrate residues P4-P1. The nitrile warhead that is the target of the nucleophilic attack by C145 is highlighted with a purple circle.

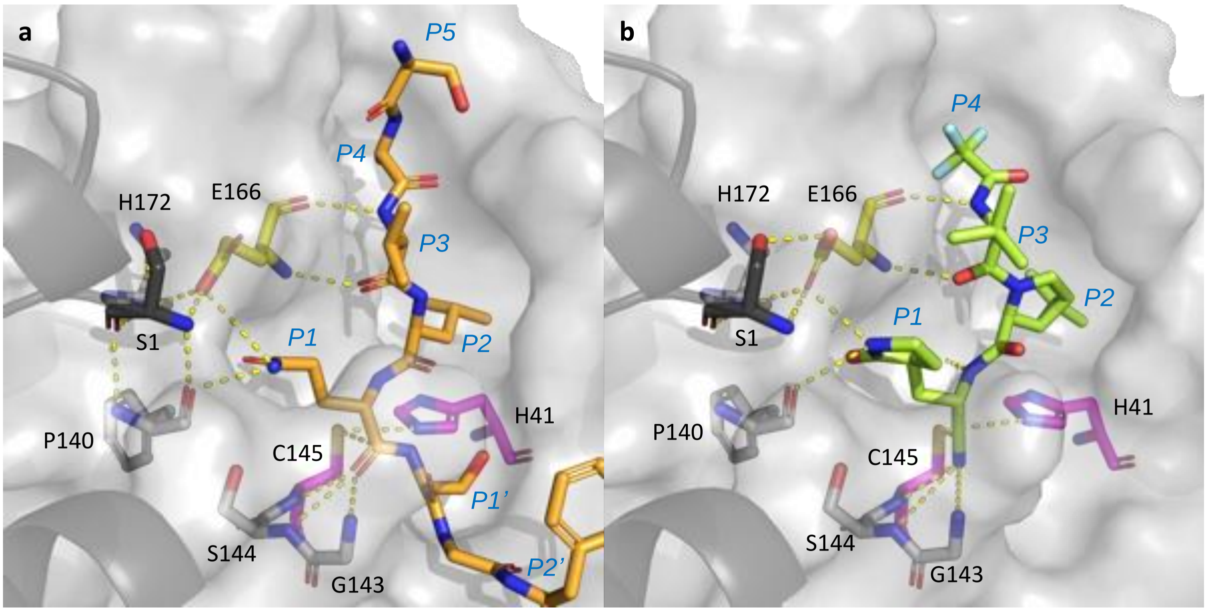


**Fig. S6. Structural basis of Mpro-substrate and Mpro-nirmatrelvir interactions.** Figure showing important Mpro residues (sticks) and interactions (yellow dashes) for catalysis and binding of **(a)** nsp4/nsp5 substrate peptide (orange sticks) and **(b)** nirmatrelvir (green sticks). E166 (yellow) stabilizes both substrate peptide and nirmatrelvir binding through three hydrogen bonds, and the Mpro dimer through interaction with S1 in the opposite monomer (black). The S1-E166 interaction is a part of a hydrogen bonding network in the S1 subsite involving, among other residues, F140 and H172. C145 in the catalytic dyad (magenta) is stabilized by a hydrogen bond to H41, and is positioned correctly for a nucleophilic attack on the substrate peptide P1 carbonyl carbon in (a) and the nirmatrelvir cyano carbon in (b). Hydrogen bonds from backbone amides of G143, S144, and C145 in the oxyanion hole to the substrate peptide P1 carbonyl oxygen in (a) and to the nirmatrelvir cyano nitrogen in (b) are shown as yellow dashes.

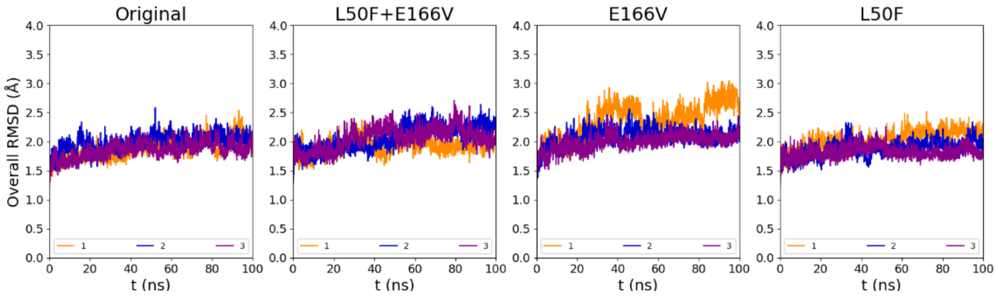


**Fig. S7. Overall RMSD for nirmatrelvir simulations calculated relative to the minimized structure.**

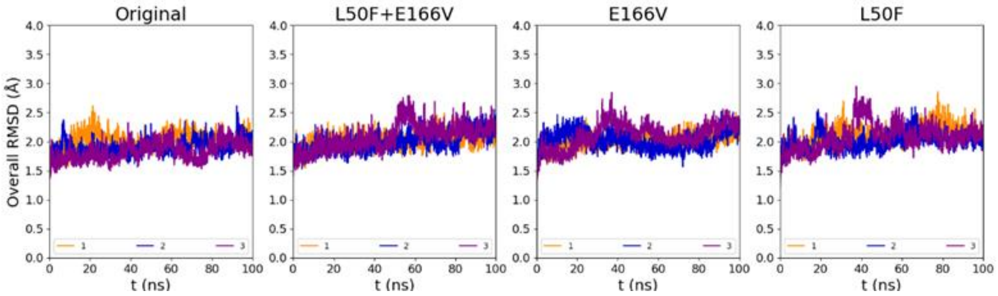


**Fig. S8. Overall RMSD for nsp4/nsp5 substrate peptide simulations calculated relative to the minimized structure.**

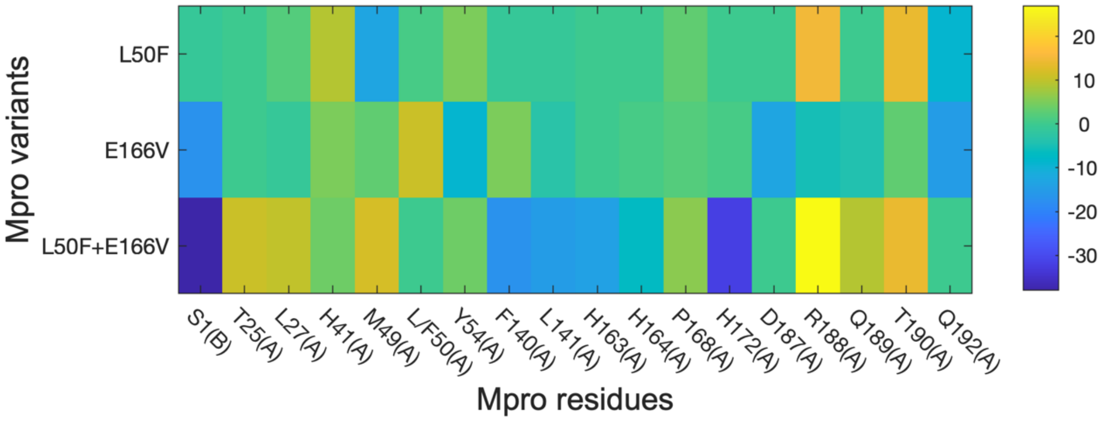


**Fig. S9. Influence of Mpro substitutions on interaction of nirmatrelvir with specific Mpro residues.** Interaction heatmap showing differences in nirmatrelvir proximities (within 3 Å) between simulations of Mpro variants and original Mpro, showing Mpro residues on the x-axis and the studied set of Mpro substitutions on the y-axis. For Mpro residues, (A) signifies location of the residue to Mpro monomer A, while (B) signifies location of the residue to Mpro monomer B. The color bar indicates increased/decreased interactions with respect to the original Mpro. Yellow and dark blue correspond to gain and reduction in interactions, respectively, for the Mpro variants when compared to the original Mpro. Thus, positive and negative percentages indicate that the occurrence of interactions is increased and decreased, respectively, in Mpro variants with substitutions compared to the original Mpro at the given Mpro residues.

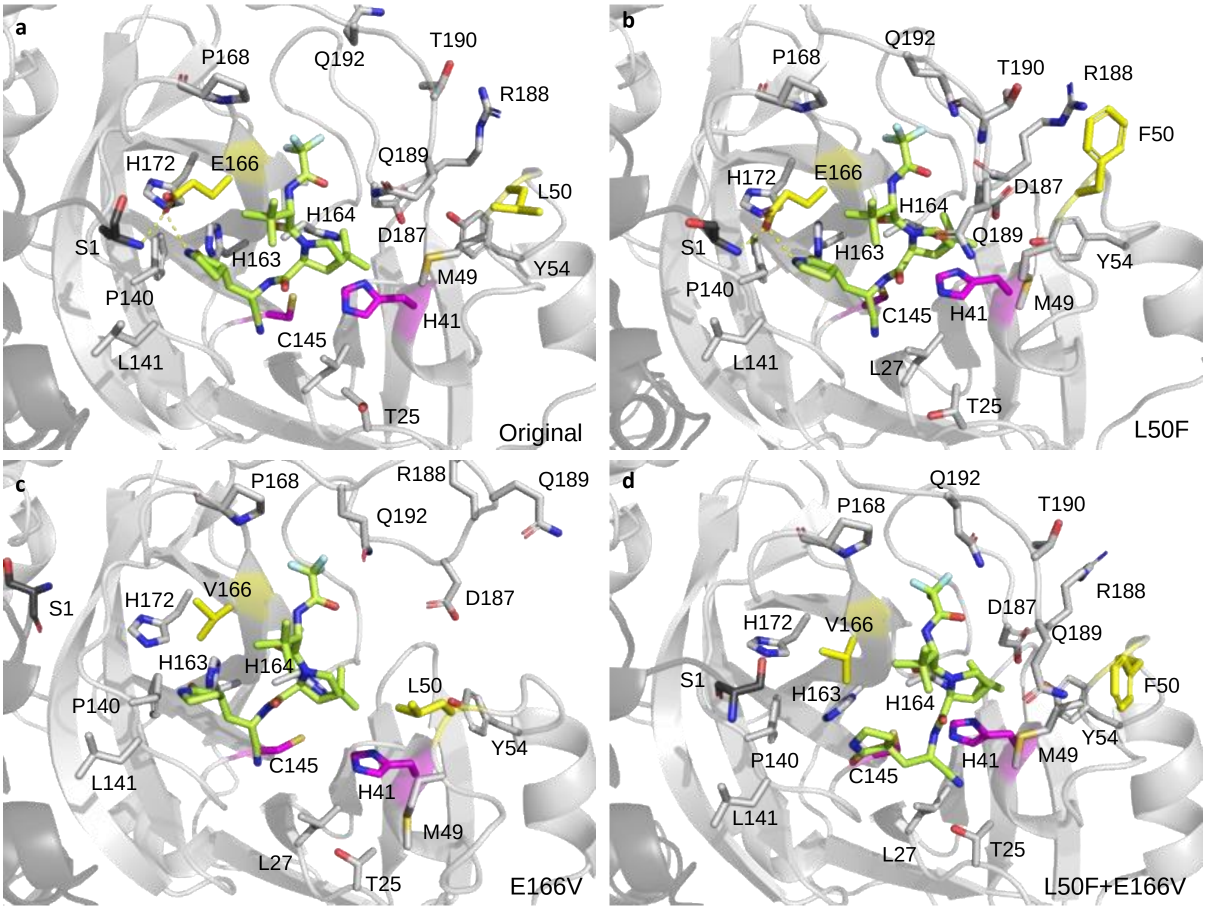


**Fig. S10. Mpro-nirmatrelvir conformations for original Mpro and Mpro with resistance associated substitutions.** Final Mpro-nirmatrelvir conformations extracted from MD simulations with labelling of Mpro residues for which changes in nirmatrelvir interactions are observed (black) for **(a)** original Mpro, **(b)** L50F, **(c)** E166V, and **(d)** L50F+E166V. The important interactions of E166 in monomer A (light grey) with S1 in monomer B (dark grey) and with nirmatrelvir (green sticks with C:green, N: blue, O:red, F:light-blue) are indicated by yellow dashes. The catalytic dyad (H41 and C145) is shown in magenta and L50F and E166V in yellow.

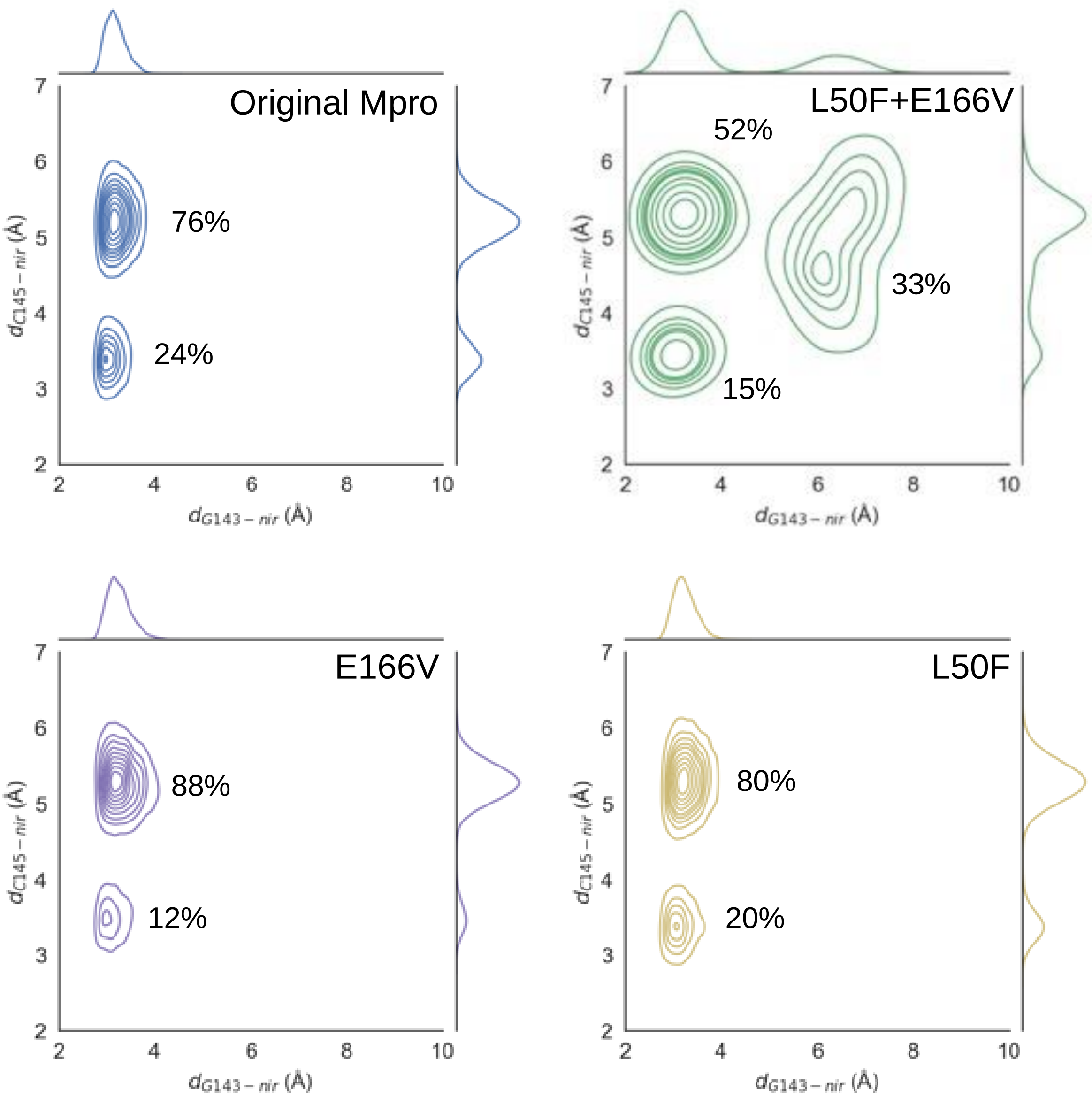


**Fig. S11. Shift of Mpro-nirmatrelvir conformational equilibrium for Mpro with resistance associated substitutions.** Contour plots with distances between C145 sulfur and nirmatrelvir cyano carbon on the y-axis and between G143 amide nitrogen and nirmatrelvir cyano nitrogen on the x-axis. For each Mpro-nirmatrelvir conformation, the frequency of the conformation is calculated and given in percentage of all simulation frames.

### Supplementary Tables

| Nucleotide change[g] | Amino acid change[g] | SARS-CoV-2 protein[h] | Escape 1[a] - %[b] 5xEC50 D16[i] | Escape 1[a] - %[b] 5xEC50 D57[i] | Escape 1[a] - %[b] 5xEC50 D74[i] | Escape 2[c] - %[b] 5xEC50 D24[i] | Escape 2[c] - %[b] 7xEC50 D57[i] | Escape 3[d] - %[b] 7xEC50 D124[i] | Escape 4[e] - %[b] 7xEC50 D24[i] | Escape 4[e] - %[b] 7xEC50 D57[i] | Escape 5[f] - %[b] 7xEC50 P3D5[i] |
|---|---|---|---|---|---|---|---|---|---|---|---|
| C10116T | T21I | Mpro | - | - | - | 16 | - | 92 | - | - | - |
| C10202T | L50F | Mpro | - | 100 | 100 | 81 | 100 | 100 | - | 100 | - |
| G10533T | C160F | Mpro | - | - | 48 | - | - | - | - | - | - |
| C10572T | A173V | Mpro | - | 98 | 100 | 81 | 100 | 100 | 98 | 100 | 78 |
| C10626T | A191V | Mpro | - | - | 20 | - | - | - | - | 100 | 14 |

**Table S1. Non-synonymous mutations in boceprevir escape viruses localizing to sequences encoding Mpro.**

[a]Escape 1, source of polyclonal escape virus BOC-EV1: continuous virus propagation in VeroE6 cells for 74 days under treatment with boceprevir. The culture was treated with 5xEC50 boceprevir from day 3 to 24 post infection, with 5xEC50 from day 28 to 36, with 4xEC50 from day 37 to 42 and with 5xEC50 from day 43 to 74. In the given time intervals, treatment was carried out every 2-3 days.

[b]Frequency (%) of non-synonymous nucleotide changes in viral genome sequence encoding Mpro recorded by next generation sequencing (NGS). Analyzed viruses were harvested from cell culture supernatant. Changes that occurred with at least 10% frequency in at least one of the analyzed virus populations were included in this table. -, frequency of the given change was <10%.

[c]Escape 2, source of polyclonal escape virus BOC-EV2: continuous virus propagation in VeroE6 cells for 57 days under treatment with boceprevir. The culture was treated with 5xEC50 boceprevir from day 2 to 53 post infection and with 7xEC50 from day 54 to 57. In the given time intervals, treatment was carried out every 2-3 days.

[d]Escape 3: continuous virus propagation in VeroE6 cells for 124 days under treatment with boceprevir. The culture was treated directly after infection with 2.5xEC50 boceprevir for 21 days, with 3xEC50 from day 22 to 27, with 4xEC50 from day 28 to 33, with 5xEC50 from day 34 to 100 and with 7xEC50 from day 101 to 124. In the given time intervals, treatment was carried out every 2-3 days.

[e]Escape 4: continuous virus propagation in VeroE6 cells for 57 days under treatment with boceprevir. The culture was treated with 7xEC50 boceprevir from day 2 to 57 post infection. In the given time interval, treatment was carried out every 2-3 days.

[f]Escape 5: primary escape culture followed by 3 viral passages in VeroE6 cells under treatment with boceprevir. All cultures were treated directly after infection and then every 2-3 days. The primary escape culture was treated with 2.5xEC50 boceprevir for 21 days, with 3xEC50 from day 22 to 27

and with 4xEC50 from day 28 to 34. The 1st passage (P1) culture was treated with 5xEC50 for 3 days. The 2nd passage (P2) culture was treated with 7xEC50 for 3 days. The 3rd passage (P3) culture was treated with 7xEC50 for 5 days.

[g]Nucleotide / amino acid position numbers and original nucleotides / amino acids, given in front of the position numbers, relate to the nucleotide / amino acid sequence of the SARS-CoV-2/human/Denmark/DK-AHH1/2020 strain (GenBank accession number MZ049597). The changed nucleotides / amino acids, acquired during continuous virus propagation or passage under boceprevir treatment, are given after the position numbers.

[h]SARS-CoV-2 protein, to which the identified amino acid changes located relating to the SARS-CoV-2/human/Denmark/DK-AHH1/2020 strain (GenBank accession number MZ049597).

[i]Specification of conditions under which viral genomes subjected to NGS were sampled. Fold EC50 applied and passage (P) and/or day (D) postinfection at sampling time.

| Nucleotide change[d] | Amino acid change[d] | SARS-CoV-2 protein[e] | Escape 1[a] - %[b] 6.25xEC50 P1D11[f] | Escape 1[a] - %[b] 12.5xEC50 P4D8[f] | Escape 1[a] - %[b] 40xEC50 P5D10[f] | Escape 2[c] - %[b] 6.25xEC50 D17[f] | Escape 2[c] - %[b] 20xEC50 P3D6[f] | Escape 2[c] - %[b] 60xEC50 P4D8[f] | Escape 2[c] - %[b] 120xEC50 P5D9[f] |
|---|---|---|---|---|---|---|---|---|---|
| T834G | F10C | nsp2 | - | 99 | 99 | - | - | - | - |
| T2827G | N36K | nsp3 | - | - | 22 | - | - | - | - |
| C10116T | T21I | Mpro | - | 96 | 100 | - | - | - | - |
| A10197G | D48G | Mpro | - | - | - | 81 | - | - | - |
| C10202T | L50F | Mpro | - | - | - | - | 31 | 98 | 99 |
| A10551T | E166V | Mpro | - | - | - | - | 100 | 99 | 99 |
| C10965T | T304I | Mpro | 99 | 100 | 100 | - | - | - | - |
| C11379T | A136V | nsp6 | - | - | - | 82 | 100 | 100 | 100 |
| T11522G | F184V | nsp6 | - | 99 | 99 | - | - | - | - |
| A12497T | N136Y | nsp8 | - | - | - | - | - | 23 | 11 |
| C13140T | T39I | nsp10 | - | 20 | - | - | - | - | - |
| A15671C | E744A | nsp12 | - | - | - | - | 56 | - | - |
| T17363C | I376T | nsp13 | - | - | - | - | 74 | 94 | 96 |
| C19547T | S503L | nsp14 | - | - | - | - | 15 | 93 | 95 |
| T24469G | N969K | S | - | - | - | - | - | 81 | 96 |
| T26300A | L19H | E | - | - | - | - | 62 | 12 | 10 |
| T26360G | L39W | E | - | - | - | - | 20 | - | - |
| T27346C | Y49H | ORF6 | - | - | - | - | - | - | 11 |
| A27862G | D36G | ORF7b | - | - | - | - | - | 36 | 51 |
| C28697T | P142S | N | - | - | - | - | - | 36 | 51 |

**Table S2. Non-synonymous mutations in nirmatrelvir escape viruses in the complete open reading frame (ORF).**

[a]Escape 1, source of polyclonal escape virus NIR-EV1: primary escape culture followed by 5 viral passages in VeroE6 cells under treatment with nirmatrelvir. All cultures were treated directly after infection and then every 2-3 days. The primary escape culture was treated with 5xEC50 nirmatrelvir for 7 days. The 1st passage (P1) culture was treated with 6.25xEC50 for 11 days. The 2nd passage (P2) culture was treated with 7.5xEC50 for 5 days. The 3rd passage (P3) culture was treated with 9xEC50 for 3 days. The 4th passage (P4) culture was treated with 20xEC50 for 8 days. The 5th passage culture (P5) was treated with 40xEC50 for 10 days.

[b] Frequency (%) of non-synonymous nucleotide changes in the complete ORF recorded by NGS. Analyzed viruses were harvested from cell culture supernatant. Changes that occurred with at least 10% frequency in at least one of the analyzed virus populations were included in this table. -, frequency of the given change was <10%.

[c]Escape 2: source of polyclonal escape virus NIR-EV2: primary escape culture followed by 5 viral passages in VeroE6 cells under treatment with nirmatrelvir. All cultures were treated directly after infection and then every 2-3 days. The primary escape culture was treated with 6.25xEC50 nirmatrelvir for 30 days. The 1st passage (P1) culture was treated with 7.5xEC50 for 5 days. The 2nd passage (P2) culture was treated with 10xEC50 for 3 days. The 3rd passage (P3) culture was

treated with 20xEC50 for 6 days. The 4th passage (P4) culture was treated with 60xEC50 for 8 days. The 5th passage culture (P5) was treated with 120xEC50 for 9 days.

[d]Nucleotide / amino acid position numbers and original nucleotides / amino acids, given in front of the position numbers, relate to the nucleotide / amino acid sequence of the SARS-CoV-2/human/Denmark/DK-AHH1/2020 strain (GenBank accession number MZ049597). The changed nucleotides / amino acids, acquired during passage under nirmatrelvir treatment, are given after the position numbers.

[e]SARS-CoV-2 protein, to which the identified amino acid changes located relating to the SARS-CoV-2/human/Denmark/DK-AHH1/2020 strain (GenBank accession number MZ049597).

[f]Specification of conditions under which viral genomes subjected to NGS were sampled. Fold EC50 applied and passage (P) and/or day (D) postinfection at sampling time.

| Nucleotide change[b] | Amino acid change[b] | SARS-CoV-2 protein[c] | SARS-CoV-2 variant-%[a] T21I P4D2[d] | L50F P4D2[d] | E166V P4D2[d] | A173V P4D2[d] | T304I P4D2[d] | L50F+ A173V P4D2[d] | L50F+ E166V P4D2[d] | T21I+ T304I P4D2[d] |
|---|---|---|---|---|---|---|---|---|---|---|
| C5183T | P822S | nsp3 | - | - | 12 | - | - | - | - | - |
| C6504A | A1262E | nsp3 | - | - | - | - | 17 | - | - | - |
| A6750G | N1344S | nsp3 | 27 | - | - | - | - | - | - | - |
| A9492C | H313P | nsp4 | - | - | 87 | - | - | - | - | - |
| C10116T | T21I | Mpro | 100 | - | | - | - | - | - | 100 |
| C10202T | L50F | Mpro | - | 100 | 26 | - | - | 100 | 100 | - |
| A10551T | E166V | Mpro | - | - | 100 | - | - | - | 100 | - |
| C10572T | A173V | Mpro | - | - | - | 89 | - | 100 | - | - |
| C10965T | T304I | Mpro | - | - | - | - | 100 | - | - | 100 |
| C11379T | A136V | nsp6 | - | - | - | 90 | - | - | - | - |
| C19325T | P429L | nsp14 | - | - | - | 12 | - | - | - | - |
| A22296G | H245R | S | - | 28 | - | - | - | - | - | - |
| A22206G | D215G | S | - | - | - | - | - | - | - | 89 |
| C23525T | H655Y | S | - | 11 | - | - | - | - | - | - |
| G23607A | R682Q | S | - | - | - | - | 14 | 87 | - | 93 |
| G23607T | R682L | S | - | - | - | - | - | - | 15 | - |
| A23618G | S686G | S | 90 | - | - | - | - | - | - | - |
| C26261T | S6L | E | - | - | - | - | - | - | 15 | - |
| C26309A | A22D | E | - | - | - | 30 | - | - | - | - |
| A27344T | K48I | ORF6 | 89 | - | - | - | - | - | - | - |

**Table S3. Non-synonymous mutations in serially passaged SARS-CoV-2 variants in the complete ORF.**

[a] Frequency (%) of non-synonymous nucleotide changes in the complete ORF of the specified SARS-CoV-2 variants following 4 viral passages in cell culture, as recorded by NGS. Analyzed viruses were harvested from cell culture supernatant. Changes that occurred with at least 10% frequency in at least one of the analyzed virus populations were included in this table. -, frequency of the given change was <10%.

[b]Nucleotide / amino acid position numbers and original nucleotides / amino acids, given in front of the position numbers, relate to the nucleotide / amino acid sequence of the SARS-CoV-2/human/Denmark/DK-AHH1/2020 strain (GenBank accession number MZ049597). The changed nucleotides / amino acids, acquired following serial passage, are given after the position numbers.

[c]SARS-CoV-2 protein, to which the identified amino acid changes located relating to the SARS-CoV-2/human/Denmark/DK-AHH1/2020 strain (GenBank accession number MZ049597).

[d]Specification of the SARS-CoV-2 variant and the time when viral genomes subjected to NGS were sampled, being passage 4 (P4) day 2 (D2) postinfection for all variants.

| | Amino acid residue[b] | SARS-CoV-2 Mpro residue[a] | | | | |
|---|---|---|---|---|---|---|
| | | T21 | L50 | E166 | A173 | T304 |
| Number of viruses[c] (Total: 10,302,924 viruses) | A | 122 | 3 | 5 | 0 | 21 |
| | R | 0 | 20 | 7 | 79 | 1 |
| | N | 80 | 0 | 4 | 66 | 54 |
| | D | 13 | 27 | 86 | 77 | 2 |
| | C | 1 | 1 | 4 | 3 | 0 |
| | Q | 1 | 0 | 4716 | 3 | 0 |
| | E | 0 | 0 | 0 | 1 | 0 |
| | G | 0 | 2 | 17 | 3 | 0 |
| | H | 1 | 13 | 130 | 0 | 1 |
| | I | 15255 | 38 | 1 | 2 | 836 |
| | L | 3 | 0 | 3 | 13 | 1 |
| | K | 0 | 11 | 7 | 2 | 1 |
| | M | 1 | 1 | 5 | 1 | 0 |
| | F | 0 | 4370 | 4 | 2 | 0 |
| | P | 3 | 11 | 1 | 7 | 4 |
| | S | 18 | 97 | 11 | 82 | 3 |
| | T | 0 | 2 | 11 | 122 | 0 |
| | W | 0 | 0 | 0 | 4 | 0 |
| | Y | 0 | 0 | 2 | 1 | 4 |
| | V | 6 | 39 | 5 | 181 | 1 |
| | del | 13 | 26 | 70 | 62 | 1 |
| | Number of substitutions[d] | 15517 | 4650 | 5082 | 709 | 929 |

**Table S4. Naturally occurring substitutions in SARS-CoV-2 Mpro.**

[a]SARS-CoV-2 Mpro residues of interest due to identification of resistance associated substitutions in this study.

[b]Amino acid identity using one letter codes; del, deletion.

[c]For this analysis, a total of 10,302,924 SARS-CoV-2 sequences were retrived from the GISAID database on April 18th, 2022.

[d]Total number of viruses with any substitution at T21, L50, E166, A173 or T304 in Mpro.

| | Nirmatrelvir | | | | | | Substrate peptide | | | | | |
|---|---|---|---|---|---|---|---|---|---|---|---|---|
| | Coul. | | LJ | | Total | | Coul. | | LJ | | Total | |
| **Original** | -160 | ±1 | -170 | ±4 | -330 | ±3 | -227 | ±14 | -239 | ±6 | -466 | ±18 |
| **L50F** | -163 | ±2 | -180 | ±8 | -343 | ±6 | -261 | ±17 | -234 | ±12 | -495 | ±24 |
| **E166V** | -147 | ±2 | -170 | ±4 | -317 | ±6 | -241 | ±12 | -250 | ±5 | -491 | ±15 |
| **L50F+E166V** | -138 | ±4 | -175 | ±5 | -313 | ±5 | -232 | ±8 | -252 | ±7 | -484 | ±13 |

**Table S5.** Overview of interaction energies extracted from the MD simulations for original, L50F, E166V, and L50F+E166V Mpro given in kJ/mol with nirmatrelvir or nsp4/nsp5 substrate peptide bound, respectively. Total interaction energies are given as well as coulombic (Coul.) and Lennard-Jones (LJ) contributions. The energies given in the table are averages of three replicates, and the uncertainties are standard errors of mean.
